# Kinetic Insights into Photoinduced Monomer–Dimer Conversion and Activation of Orange Carotenoid Protein

**DOI:** 10.1101/2025.05.26.656073

**Authors:** Tadayuki Tokashiki, Takatoshi Ohata, Shunrou Tokonami, Takashi Oda, Yusuke Nakasone, Masahide Terazima

## Abstract

The Orange Carotenoid Protein (OCP) is a blue-green light sensor that regulates non-photochemical quenching (NPQ) in cyanobacteria through reversible transitions between its dark-adapted (OCP^O^) and light-adapted (OCP^R^) states. Despite extensive studies, the detailed reaction scheme remains unclear. In this study, we examined the photo-induced reaction dynamics of OCP using size-exclusion chromatography (SEC), small-angle X-ray scattering (SAXS), and transient grating (TG) spectroscopy. We found that OCP^O^ and OCP^R^ exist in monomer–dimer equilibria, with OCP^R^ forming more stable and elongated dimers. TG measurements revealed that upon photoexcitation, OCP^O^ monomers undergo two structural transitions before associating into OCP^R^ dimers. In contrast, OCP^O^ dimers dissociate prior to the structural rearrangement, highlighting a fundamental difference in their reaction pathways. Moreover, dimerization was found to moderately reduce the photo-reactivity of OCP^O^ compared to the monomer. We also found that apo-OCP readily forms heterodimers with OCP^R^, potentially altering reaction pathways and masking true kinetic behavior.

## 1. Introduction

Photosynthesis is a fundamental biological process that sustains life by converting solar energy into chemical energy. While light is essential for this reaction, excessive illumination can damage the photosynthetic reaction centers.^1^ To prevent such photodamage while maintaining efficient light harvesting, photosynthetic organisms have developed regulatory mechanisms such as non-photochemical quenching (NPQ).^2,3^ In cyanobacteria, NPQ is mediated by the Orange Carotenoid Protein (OCP), a 35 kDa soluble blue-green light sensor protein that binds a carotenoid chromophore.^4,5^

OCP consists of two domains: the N-terminal domain (NTD, residues 1–165), primarily consisting of α-helices, and the C-terminal domain (CTD, residues 190–317), composed of α-helices and β-sheets. These domains are connected by a flexible 25-residue linker. Additionally, the N-terminal extension (NTE, residues 1–20) associates with the CTD surface in the dark-adapted form. In the dark-adapted state, referred to as OCP^O^, a carotenoid molecule spans the two domains, interacting via π-π interactions with residues Y44 and W101 in the NTD and forming hydrogen bonds with Y201 and W288 in the CTD.^6,7^ Upon photoexcitation by blue-green light, the carotenoid undergoes isomerization and shifts approximately 12 Å toward the NTD.^8,9^ This movement breaks the hydrogen bonds^10^ and disrupts the interdomain interaction, initiating a sequence of conformational changes that transform OCP^O^ into the red-shifted, light-adapted state known as OCP^R^.^8,9,11–15^

The photoreaction of OCP involves multiple intermediate states and structural transitions occurring over a broad timescale ranging from picoseconds to hundreds of milliseconds. Spectroscopic studies using transient absorption, fluorescence, FTIR, and Raman spectroscopy have identified several intermediates and proposed mechanisms involving carotenoid translocation, NTE dissociation, and domain separation.^8–10,13–22^ However, the proposed reaction schemes vary considerably, and a unified model has not been established.

One reason for the inconsistency is the oligomerization behavior of OCP. Although OCP^O^ was long assumed to be monomeric in solution, all known crystal structures show dimer formation^7^, and recent SAXS and MS studies have confirmed that both OCP^O^ and OCP^R^ exist predominantly as dimers in solution.^23–25^ Cryo-EM studies of the OCP^R^–PBS complex further suggest that dimerization facilitates OCP–PBS interaction and stabilization.^26^

Another complicating factor is the presence of apo-OCP, a carotenoid-free form that can be co-purified during recombinant expression. Apo-OCP lacks the bridging carotenoid and therefore adopts a more open conformation resembling OCP^R^.^22,27^ The coexistence of apo-OCP may lead to unexpected photoreaction dynamics, including formation of heterodimers with photoactivated OCP. Thus, rigorous sample preparation and verification are essential for accurate kinetic and structural studies.

Although structural information on OCP has accumulated in recent years, time-resolved studies that directly track the progression of conformational rearrangements and oligomerization events remain limited. In this study, we focused on elucidating the light-dependent structural dynamics of OCP of tag-free, canthaxanthin-binding OCP by combining multiple complementary techniques. The oligomeric states under dark and light conditions were assessed using SEC and SAXS, while transient grating (TG) spectroscopy^28–32^ was used to monitor changes in molecular diffusion that reflect structural and oligomeric dynamics with high temporal resolution. To isolate the contribution of monomeric species, we also examined a monomeric mutant (R27L)^33^, which has been shown to predominantly exist as a monomer in the dark state.

This integrated approach allowed us to explore the dynamic features of the OCP photoreaction, with particular attention to the behavior of different oligomeric forms and the impact of apo-OCP. The findings contribute to a deeper understanding of the mechanisms underlying OCP function and its role in modulating NPQ activity through light-dependent structural changes.

## 2. Materials and methods

### 2.1 Protein expression and purification

OCP from *Synechocystis sp.* PCC 6803 (slr1963) was expressed in *Escherichia coli* using a previously reported co-expression system with carotenoid biosynthesis enzymes.^34^ The expression plasmid pET-Duet was used for OCP and CrtW (a ketolase for converting β-carotene to canthaxanthin), while pAC-BETA was used for β-carotene production. Apo-OCP was expressed using only the pET-Duet plasmid.

Expression of the target protein was conducted with the auto-induction method.^35^ After overnight cultivation, cells were harvested by centrifugation and resuspended in PBS buffer (137 mM NaCl, 8.1 mM Na₂HPO₄, 2.68 mM KCl, 1.47 mM KH₂PO₄, pH 7.5). After addition of DNase and RNase, the suspension was sonicated and centrifuged. Ammonium sulfate was added to the supernatant to 30% saturation to remove impurities, followed by precipitation of OCP at 60% saturation. The pellet was dissolved in 20 mM Tris-HCl (pH 8.0), and salts were removed via a desalting column. Further purification was performed using anion exchange chromatography (HiTrap Q, Cytiva), hydrophobic interaction chromatography (HiTrap Butyl HP, Cytiva), and size-exclusion chromatography (Superdex 200 Increase 10/300 GL, Cytiva). SDS-PAGE was used to confirm purity. Samples with an absorbance ratio (A_476_/A_280_) > 2.3 were used as holo-OCP. Apo-OCP was prepared similarly. The R27L mutant was generated by site-directed mutagenesis using appropriate primers and confirmed by sequencing. Protein concentrations were determined by absorbance at 476 nm (ε = 118,000 M^−1^cm^−1^ for holo-OCP) or 280 nm (ε = 35,000 M^−1^cm^−1^ for apo-OCP).

### 2.2 Absorption spectra and time-dependent absorbance measurements

Absorption spectra were measured using a diode array spectrophotometer (Agilent Cary 8454). Time-dependent absorbance changes at 550 nm were monitored under blue LED illumination (460 nm, CHR-3S, Nissin Electronic Co.) at a 60° incident angle. The path length was 1 mm and the temperature was maintained constant at 25 ℃.

### 2.3 Size-Exclusion Chromatography (SEC)

SEC measurements were conducted using Superdex 200 Increase 10/300 GL or 3.2/300 columns (Cytiva) equilibrated with PBS buffer. Absorbance at 280, 476, and 620 nm was monitored. For OCP^O^, columns were wrapped in aluminum foil; for OCP^R^, samples were continuously illuminated with a Xe lamp (Max302, Asahi Spectra) during elution. Molecular weights were estimated using a gel filtration standard (Merck). All experiments were performed at 4 °C.

### 2.4 Small Angle X-ray Scattering (SAXS)

SAXS measurements were performed at beamline BL15A2 of the Photon Factory. Samples were flowed at 0.02 mL/min to minimize radiation damage. For OCP^R^ measurements, samples were illuminated with a 450 nm laser (L450P1600MM, Thorlabs). Scattered X-rays were recorded with a PILATUS2M detector positioned 1,000 mm from the sample. The scattering vector *q* = 4πsinθ/λ (λ = 1.0 Å) was used, and data were processed using the ATSAS package.^36,37^ *R*_g_ and *I*₀ were determined from Guinier plots, and *D*_max_ and *P*(r) from GNOM.^38^ Ab initio models were reconstructed using GASBOR.^39^ Simultaneous absorption spectra were recorded using a white light source and spectrometer (Flame, Ocean Insight) to confirm the OCP^R^ state.

Porod volumes of proteins in each solution was calculated on the observed SAXS profiles to estimate the molecular weight in the solution.^40^ The Porod volume *V*_p_ corresponds to the volume of the protein in solution, and by multiplying this volume by the protein density *ρ*_protein_ and Avogadro constant *N*_A_, the molecular mass *M* of the protein in solution can be determined. The volume *V*_p_ of the solute molecule can be determined by the following equation using the relatively wide-angle region (*q*≤ 0.5 Å^−1^) of the SAXS profile with *I*_0_ obtained from the Guinier analysis.

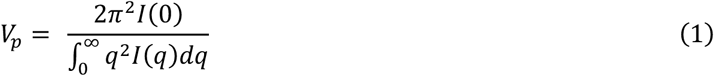

In this study, the value *ρ*_protein_=1.34 g/cm^3^ was used as the density of OCP, based on structural information computed with SOMO^41^, and the molecular mass was calculated using:

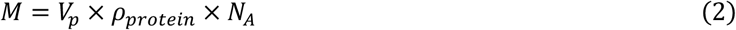

### 2.5 Transient Grating measurement

Experimental setup for the TG measurement is similar to that previously reported.^28,42–46^ The sample solution was irradiated by a pulsed light from a dye laser (480 nm, HyperDye 300, Lumonics) pumped by the third-harmonic light of a Nd:YAG laser (355 nm, Nanosecond Dye Laser: ND6000, Continuum). A diode laser (785 nm, L785P090, Thorlabs, NJ, USA) was used as the probe laser. The TG signal was detected by a photomultiplier tube (R1477, Hamamatsu Photonics, Japan) and recorded with a digital oscilloscope (DSOS054A, Agilent technologies, CA, USA). The grating wavenumber *q* for each measurement was determined from the decay rate of the thermal grating signal of an aqueous solution of bromocresol purple. To avoid re-excitation of photoproducts or reaction intermediates, the sample was gently stirred after each laser pulse, except during successive pulse excitation experiments where multiple pulses were intentionally applied without stirring to study accumulation effects.

## 3. Results

### 3.1 Oligomeric state of OCP characterized by SEC

The oligomeric states of OCP in the dark and light states were measured by SEC. Fig.2(a) depicts the SEC profiles under the dark and continuous light-illuminated conditions at an injection concentration of 100 *μ*M. The molecular mass of the elution peak upon light illumination (73 kDa) is larger than that in the dark-adapted state (37 kDa). Given that the calculated molecular mass of OCP monomer is 35 kDa, OCP^O^ exists mainly as a monomer, while OCP^R^ is predominantly dimeric at this concentration.

**Figure 1.**
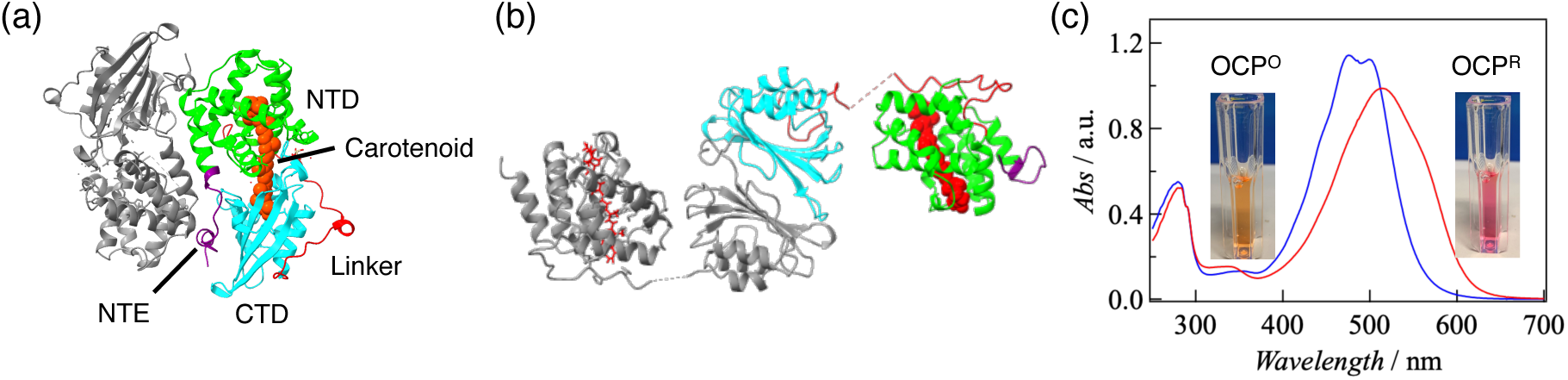
(a) The structure of the OCP^O^ dimer (PDB ID: 3MG1). (b) The structure of the OCP^R^ dimer in complex with the phycobilisome (PBS) (PDB ID: 7SC9). In the OCP^O^ dimer, each protomer interacts through its N-terminal domain (NTD) and the C-terminal domain (CTD) of the opposing protomer, whereas in the OCP^R^ dimer, the two monomers associate via the surfaces of their CTDs. (c) Absorption spectra of OCP^O^ and OCP^R^, with inset images showing the apparent color differences between the two states.

**Figure 2.**
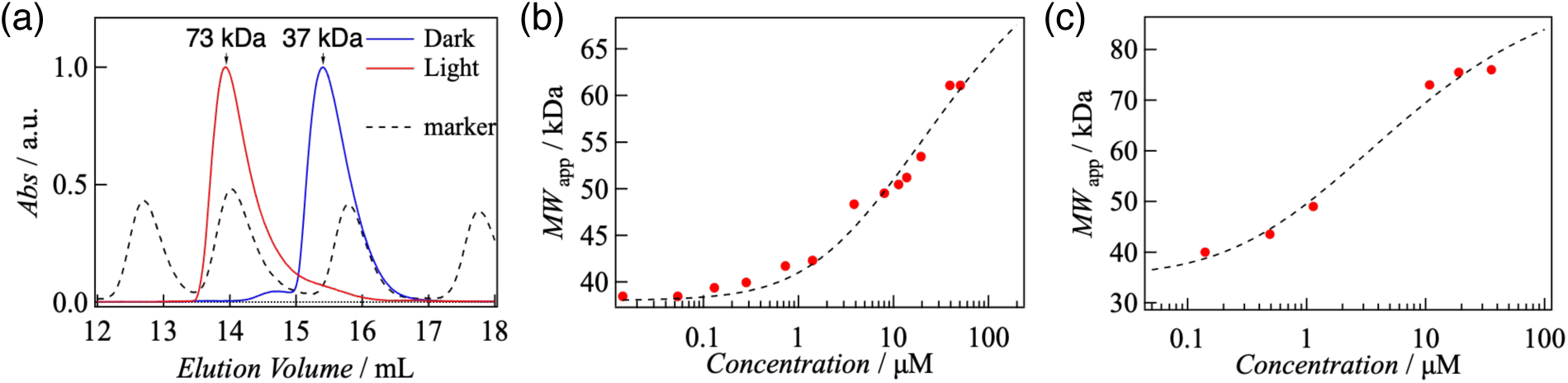
(a) SEC profiles of OCP in the dark-adapted state (blue) and under Xe lamp illumination (red), monitored at 280 nm and normalized to the elution peak intensity. Dotted lines indicate molecular weight markers, with corresponding elution volumes of 142 kDa, 67 kDa, 32 kDa, and 1.4 kDa from left to right. (b, c) Relationship between apparent molecular weight and concentration at the elution peak in the dark-adapted state (b) and under illumination (c).

To evaluate the concentration dependence of oligomerization, SEC measurements were performed at various concentrations (Figure S1). The molecular masses estimated from the elution peaks were plotted against the concentration calculated from absorbance at the detection point (Fig. 2(b) and (c)). In both dark and light states, the molecular mass increased with concentration, indicating monomer–dimer equilibria.

The dissociation constants in the dark- and light-adapted states are determined from these plots. The apparent molecular mass (<M>) in the monomer-dimer equilibrium may be given by Eq. (3).

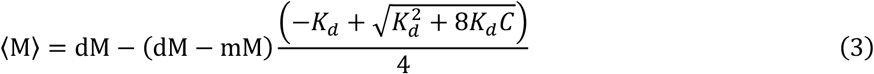

where dM and mM are the molecular masses of the dimer and monomer, *K*_d_ is the dissociation constant, and *C* is the total OCP concentration. The concentration dependence of <M> (Fig.2(b) and (c)) are fitted by this equation to determine mM, dM, and *K*_d_ in the dark state (mM_D_, dM_D_, and *K*_dD_), and the light state (mM_L_, dM_L_, and *K*_dL_). From this analysis, the fitted parameters were determined as follows: mM_D_ = 38 kDa, dM_D_ = 78 kDa, *K*_dD_ = 10 µM (dark state); mM_L_ = 35 kDa, dM_L_ = 94 kDa, *K*_dL_ = 1.8 µM (light state). The differences between mM and dM values in dark and light conditions may reflect conformational changes, which are further discussed in the SAXS section. It is also important to note that these *K*_d_ values may be underestimates, as dilution occurs in the column and the effective concentration at the detection point may be lower.

To assess whether heterodimers form between OCP^O^ and OCP^R^, we conducted SEC measurements under partially activating light intensity, where absorption spectra confirmed coexistence of both states. At 100 µM (the injected concentration), the SEC profile showed two peaks corresponding to monomer and dimer (Fig. S2). The absorbance ratios at 280/620 nm (A280/A620) were 5.3 for the dimer peak and 24 for the monomer peak. Given that A280/A620 is ∼30 in the dark and ∼5.8 in the light (Fig. 1(c)), this result indicates that the dimer peak mostly consists of OCP^R^ dimers, while the monomer peak is primarily OCP^O^ monomer with a minor fraction of OCP^R^ monomer. These results suggest that heterodimers between OCP^O^ and OCP^R^ are not formed under these conditions.

We further analyzed the oligomeric state of the OCP(R27L) mutant (Fig. S3), which is known to remain monomeric in the dark^33^. SEC profiles confirmed that even at concentrations up to 200 µM, the elution peak position remained unchanged in the dark, consistent with stable monomeric behavior. However, under light illumination, a concentration-dependent shift toward higher molecular mass was observed, indicating that OCP(R27L) adopts a monomer–dimer equilibrium in the light state.

### 3.2 Solution structure of OCP elucidated by SAXS

SAXS measurements were conducted to investigate the solution structures of OCP^O^ and OCP^R^ dimers. At a protein concentration of 100 µM, the scattering profiles of OCP changed upon light illumination, indicating light-induced conformational changes (Fig. 3(a)). Guinier analysis in the small-angle region showed that the forward scattering intensities (I₀) under dark and illuminated conditions were nearly identical (*I*₀ = 0.13), and Porod volume analysis^40^ revealed molecular masses of 72 kDa for the dark state and 77 kDa for the light-adapted state, consistent with dimeric forms of both OCP^O^ and OCP^R^ at this concentration.

**Figure 3.**
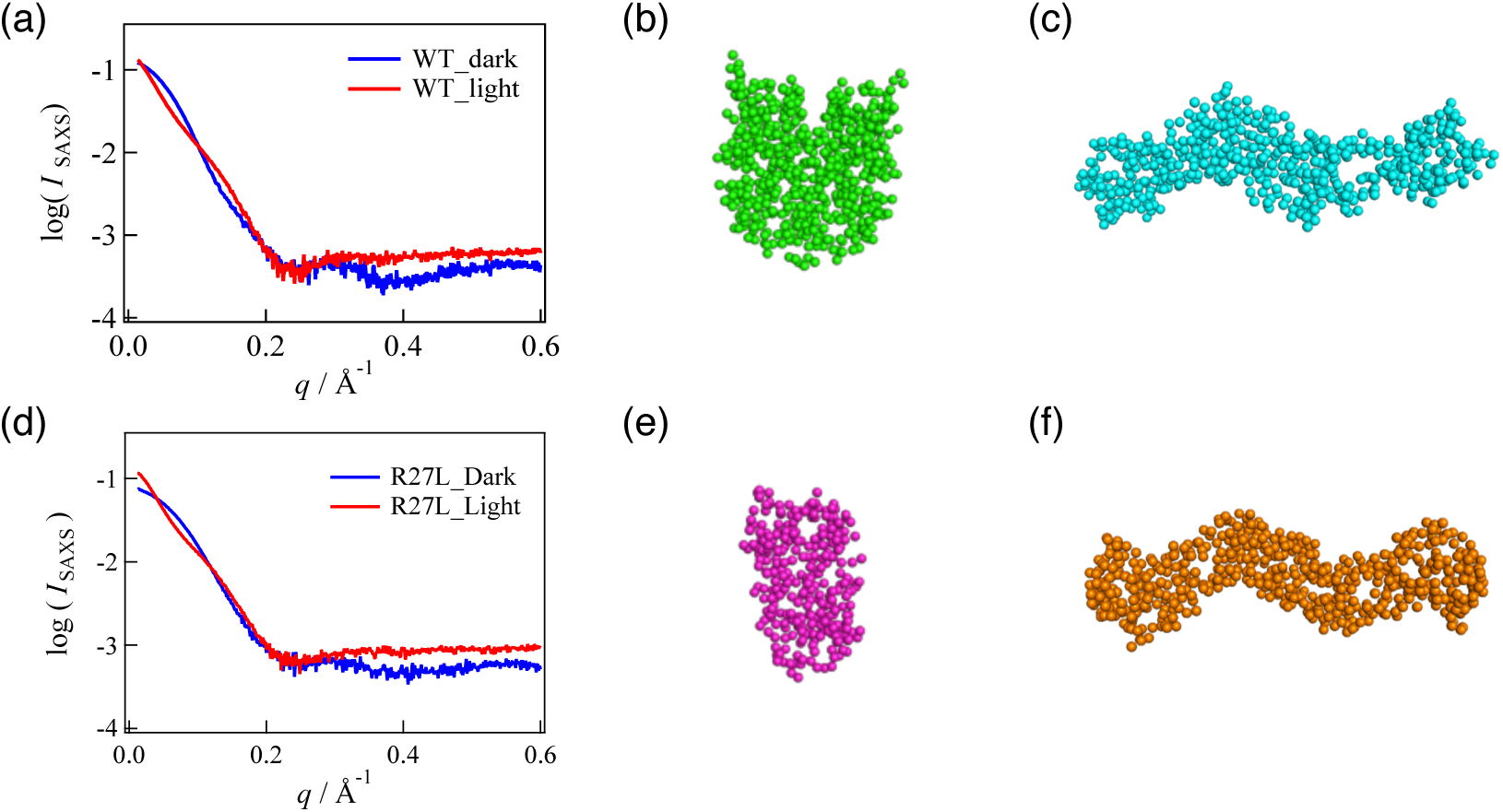
(a) SAXS profiles of WT-OCP in the dark-adapted state (blue line) and light-adapted state (red line) at a concentration of 100 μM. (b, c) Predicted solution structures of wild-type OCP in the dark-adapted state (b) and light-adapted state (c). (d) SAXS profiles of OCP(R27L) in the dark-adapted state (blue line) and light-adapted state (red line) at a concentration of 100 μM. (e, f) Predicted solution structures of OCP(R27L) in the dark-adapted state (e) and light-adapted state (f).

In contrast, the radius of gyration (*R*_g_) increased markedly from 26.6 Å in the dark to 38.3 Å under illumination (Fig. S4(a)(b)). Theoretical *R*_g_ values for monomeric and dimeric forms of OCP^O^ and OCP^R^ were calculated using SOMO^41^ based on high-resolution crystal and cryo-EM structures (PDB ID: 3MG1 and 7SC9), and are summarized in Table 1. The experimentally observed *R*_g_ values agree well with those calculated for the dimeric structures, supporting the conclusion that both OCP^O^ and OCP^R^ exist as dimers under these conditions. Radial distribution function analysis using GNOM^38^ further showed that the maximum particle dimension (*D*_max_) increased from 89.6 Å in the dark to 137 Å in the light (Fig. S4(c)). The significant changes in *R*g and *D*_max_ suggest a drastic conformational change upon photoactivation, likely reflecting domain dissociation between the NTD and CTD and resulting in an elongated structure for OCP^R^. Ab initio shape reconstructions using GASBOR^39^ based on the scattering profiles (Fig. 3(b)(c)) supported this interpretation. These models indicated that dimeric OCP^O^ adopt compact conformations, whereas the OCP^R^ dimer exhibits an extended shape with increased separation between domains.

**Table 1.** Structural parameters of WT and R27L mutant obtained from SAXS profile analyses, along with calculated parameters for OCP^O^ and OCP^R^ using SOMO and high-resolution structures (PDB ID: 3MG1 for the dark-adapted state and 7SC9 for the light-adapted state).

|  | WT |  | R27L |  | OCP<sup>O</sup> (calculation) |  | OCP<sup>R</sup> (calculation) |  |
| --- | --- | --- | --- | --- | --- | --- | --- | --- |
|  | Dark | Light | Dark | Light | monomer | dimer | monomer | dimer |
| *MW* /kDa | 72 | 77 | 39 | 63 | 35 | 70 | 35 | 70 |
| *R<sub>g</sub>* /Å | 26.6 | 38.3 | 22.3 | 39.0 | 20.7 | 26.3 | 24.5 | 35.8 |
| *D<sub>max</sub>* /Å | 89.6 | 137 | 70.4 | 128 | 69 | 91 | 70 | 125 |

The SAXS profile of the OCP(R27L) mutant is shown in Fig. 3(d). Upon light illumination, both the profile and forward scattering intensity changed. Guinier analysis (Fig. S4(d)(e)) and Porod volume analysis indicated an increase in *R*_g_ from 22.3 Å to 39.0 Å and in molecular mass from 39 kDa to 63 kDa. These findings are consistent with SEC data showing that OCP(R27L) exists as a monomer in the dark and forms dimers upon light exposure. The D_max_ values calculated from the *P*(r) distribution were 70.4 Å in the dark and 128 Å in the light (Fig. S4(f)). The ab initio models (Fig. 3(e)(f)) reconstructed from the scattering profiles using GASBOR are consistent with these results, showing a compact monomeric structure in the dark state and an elongated dimeric structure with increased domain separation in the light-adapted state.

### 3.3 Photoinduced dimerization of OCP probed by TG method

The photoreaction dynamics of OCP, driven by pulsed light excitation, were investigated using TG method. Figure 4(a) shows the TG signal at *q*^2^ = 5.9 × 10^9^ m^−2^ and a protein concentration of 100 µM. Immediately after excitation, a strong signal appears and decays within milliseconds. Following this, a much weaker rise-and-decay signal is observed (Fig. 4(a), inset). In general, the time-profile of the TG signal is expressed by the following equation (Eq. (4)).^30,32^

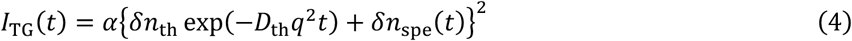

**Figure 4.**
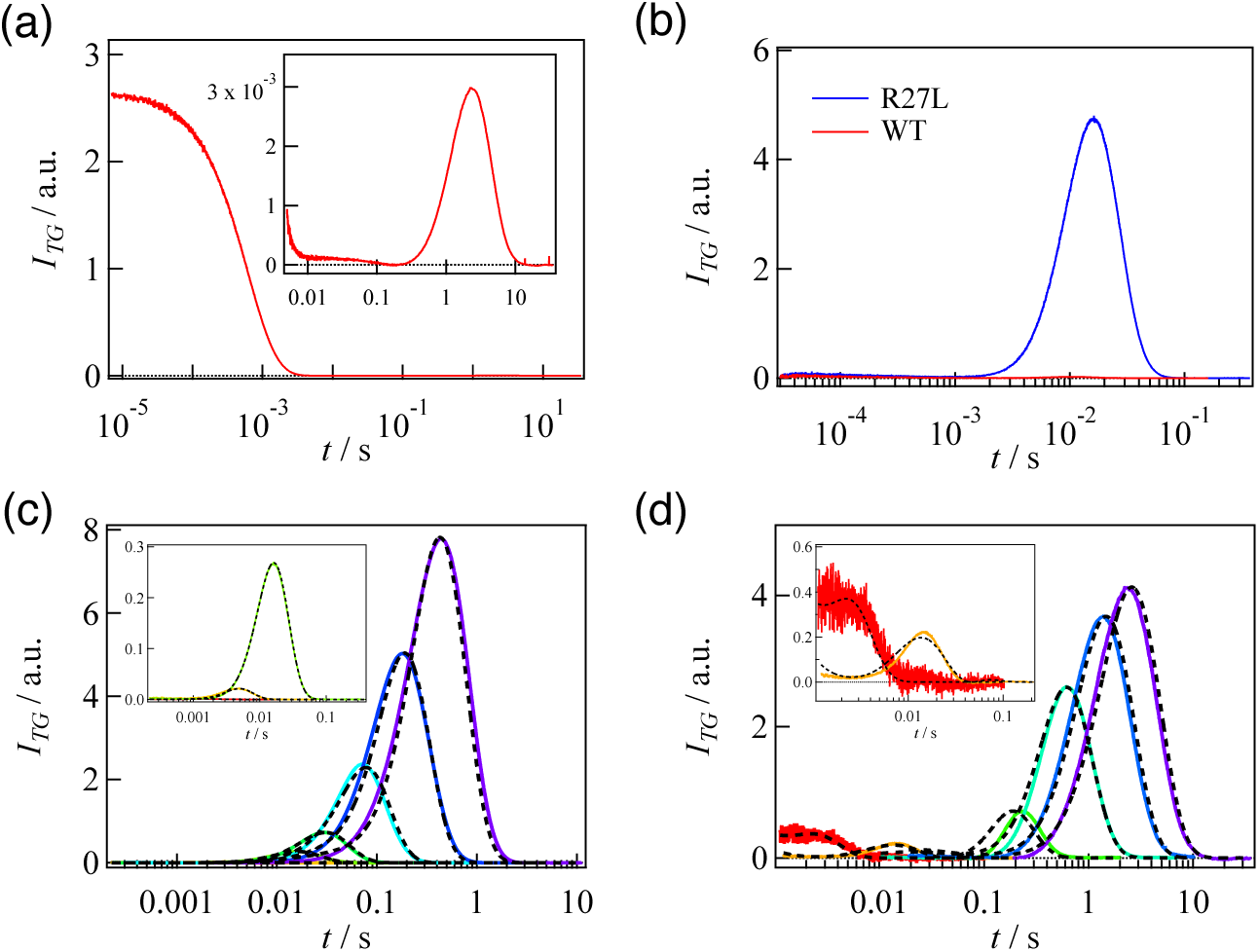
(a) TG signal of WT-OCP at a grating wavenumber of *q*^2^ = 5.9 × 10^9^ m^−2^ with a sample concentration of 100 μM. The inset shows an expanded view of the molecular diffusion signal. (b) Comparison of TG signals for WT-OCP (blue) and the R27L mutant (red) at 100 μM and *q*^2^ = 1.3 × 10^12^ m^−2^. The molecular diffusion signal was approximately 200 times stronger for the R27L mutant. (c) Grating wavenumber dependence of TG signals for OCP(R27L) (grating wavenumbers: red, 1.7 × 10^13^ m^−2^; orange, 4.8 × 10^12^ m^−2^; light green, 1.3 × 10^12^ m^−2^; green, 6.4 × 10^11^ m^−2^; cyan, 2.4 × 10^11^ m^−2^; blue, 9.0 × 10^10^ m^−2^; purple, 3.7 × 10^10^ m^−2^). Dotted lines show the fitting curves based on Scheme 1. The inset shows an expanded view of the early time region. (d) Grating wavenumber dependence of TG signals for WT-OCP (grating wavenumbers: red, 7.3 × 10^12^ m^−2^; orange, 6.1 × 10^11^ m^−2^; light green, 1.2 × 10^11^ m^−2^; green, 2.9 × 10^10^ m^−2^; blue, 1.1 × 10^10^ m^−2^; purple, 5.9 × 10^9^ m^−2^). Dotted lines show the fitting curves based on Scheme 3. The inset shows an expanded view of the early time region.

where α is a constant representing the system sensitivity, δ*n*_th_ is the pre-exponential factor of the thermal grating signal, *D*_th_ is the thermal diffusivity of the solution, and δ*n*_spe_(t) represents the change in the refractive index due to changes in the optical absorbance and molecular volumes. The fast-decaying component is assigned to thermal grating generated by nonradiative deactivation of excited carotenoids, while the slower rise-and-decay component is attributed to molecular diffusion signal since its time-range depends on *q* (Figure. 4(c)). ^31,32,47^

The weak intensity of the diffusion signal indicates a low photoreaction efficiency. Since δ*n*_th_ is negative under the present conditions, the rise and decay components appear as negative and positive signals, respectively, corresponding to the diffusion of the reactant and the slower-diffusing photoproduct. Before analyzing this signal in detail, it is important to note that SEC and SAXS analyses have shown that OCP exists in a monomer–dimer equilibrium even in the dark state, requiring consideration of photoreactions originating from both species. This complexity makes interpretation of the TG signal more challenging. To simplify the analysis and specifically probe the monomer reaction, we performed TG measurements using the monomeric mutant OCP(R27L).

The TG signal of the monomeric mutant OCP(R27L) at *q*^2^ = 1.3 × 10^12^ m^−2^ and 100 µM is shown in Fig. 4(b), with the signal of wild type OCP (WT-OCP) included for comparison. OCP(R27L) exhibited thermal signal and rise-decay profile of molecular diffusion, with assignment of the rise and decay corresponding to diffusion of the reactant and the photo-product, respectively, which is consistent with the analysis of WT-OCP. A striking observation is that the diffusion signal of OCP(R27L) is approximately 200 times stronger than that of WT. This signal intensity reflects both the magnitude of *D* change and the photoreaction efficiency. As will be shown in later analysis, the *D* change associated with the monomer reaction is identical for both WT-OCP and OCP(R27L). Therefore, the enhanced diffusion signal intensity in OCP(R27L) is attributed to its higher fraction of reactive monomer. SEC and SAXS analyses have demonstrated that the main photoreaction of OCP(R27L) involves light-induced dimerization, and that WT-OCP undergoes a shift in monomer–dimer equilibrium toward the dimeric form upon illumination. These observations suggest that the observed *D*-decrease reflects an increase in molecular volume caused by photo-induced dimer formation.

Moreover, SEC analyses indicate that OCP^R^ forms stable homodimers rather than heterodimers with OCP^O^. To examine whether this homodimerization behavior also appears under TG conditions, we conducted repeated pulsed excitations at 0.1 Hz for WT-OCP. Under this condition, where the excitation interval of 10 s is much shorter than the dark recovery time (>40 s), repeated excitations progressively increase the population of OCP^R^ monomers. This accumulation enhances the diffusion signal, as newly generated OCP^R^ molecules have more opportunities to associate with surrounding OCP^R^, promoting the formation of OCP^R^-dimer. In contrast, if the diffusion signal were due to intramolecular conformational changes or heterodimer formation between OCP^R^ and OCP^O^, the signal intensity would be expected to decline with repeated excitation due to the depletion of OCP^O^.

The diffusion signal increased over successive pulses and eventually reached a plateau (Fig. S5(a)(b)). Importantly, we confirmed that the accumulated OCP^R^ species itself did not undergo further photoinduced reactions that would contribute to the diffusion signal (Fig. S5(c)). Hence, the observed increase reflects the progressive accumulation of OCP^R^ monomer, which accelerates the formation of OCP^R^-dimer and thereby enhances the intensity of the diffusion signal, in agreement with the reaction scheme established from SEC and SAXS analyses.

### 3.4 Reaction kinetics of monomer mutant OCP(R27L)

To investigate the reaction dynamics of OCP, TG signals were measured at various grating wavenumbers. We first focused on OCP(R27L) to examine the photoreaction scheme of the OCP^O^ monomer. Figure 4(c) shows the grating wavenumber (*q*) dependence of the TG signal. The diffusion signal becomes more pronounced at smaller *q* values, indicating that the photoreaction involves changes in the diffusion coefficient over timescales from milliseconds to seconds.

Since the dimerization step involves bimolecular association between two photoactivated species, the reaction– diffusion equations cannot be solved analytically, necessitating the use of numerical simulations. For these calculations, the concentration of photoactivated molecules (*C*₀) is required; however, its actual value cannot be directly determined because some molecules may undergo multiple cycles of activation and relaxation (on the picosecond timescale) within the duration of the nanosecond excitation pulse. In this study, we estimated *C*₀ to be 0.3 µM by multiplying the total protein concentration (100 µM) by the reaction quantum yield (0.3%). Numerical calculations were then performed using this estimated value.

We initially modeled the reaction as a two-step process: a rapid formation of an intermediate (I_1_) immediately following photoexcitation, followed by a direct dimerization (R → I_1_ → P). However, this model failed to reproduce the *q*-dependence of the TG signals. To improve the fit, we added a second intermediate (I_2_), creating a three-step model (R → I_1_ → I_2_ → P), but this still did not capture the early-time signals observed in the data. Only after introducing a third state (I_3_), resulting in a four-step model (Scheme 1), were we able to accurately reproduce the full TG signal profile. The subscript “m” is used to indicate that the labeled species and rate constants arises from the photoactivation of the OCP^O^ monomer, in order to distinguish these from those originating from the OCP^O^ dimer, which will be analyzed later. The superscripts “M” and “D” specify the oligomeric state (monomer or dimer) of each species at the corresponding stage in the reaction scheme.

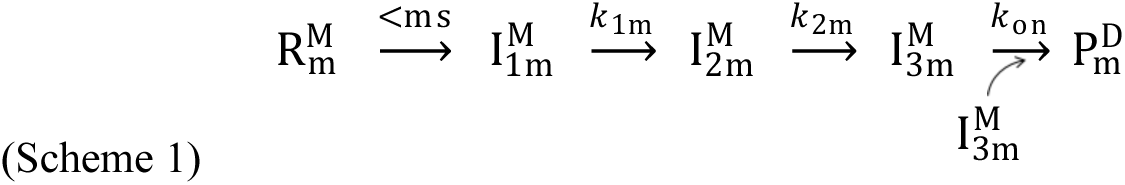

This model assumes that the final step (I_3m_ → P_m_) represents the dimerization of two OCP^R^ molecules, and that two sequential structural transitions precede this final bimolecular association step. The parameters obtained from the fitting procedure are summarized in Table 2. Specifically, numerical calculations for OCP(R27L) yielded *D*_R_ = 8.53 × 10^−1^ m^2^/s, *D*_P_ = 6.46 × 10^−1^ m^2^/s, and a bimolecular rate constant *k*_on_ = 30 µM^−1^s^−1^.The *D*_R_-value is reasonable to that expected for a protein that is the size of OCP-monomer (35 kDa); e.g., the *D* of Ovalbumin (45 kDa) is 8.7×10^−11^ m^2^/s.^48^

**Table 2.** Fitting parameters from TG analyses. Diffusion coefficients are given in units of 10^−1^ m^2^ s^−1^; the reaction rate constants for the first and second steps (*k*₁ and *k*₂) are in s^−1^. The association rate constant is given in μM^−1^ s^−1^ and was obtained assuming a total initial reactant concentration of 0.3 μM.

| | $D_R$ | $D_{I_1}$ | $D_{I_2}$ | $D_{I_3}$ | $D_P$ | $D_{P_2}$ | $k_1$ | $k_2$ | $k_{\text{on}}$ |
| --- | --- | --- | --- | --- | --- | --- | --- | --- | --- |
| **R27L** | 8.53 | 8.53 | 8.51 | 7.79 | 6.46 | — | $1.0 \times 10^3$ | 88 | 30 |
| **WT** | 7.66 | 7.66 | 8.51 | 7.79 | 6.46 | 8.53 | $1.0 \times 10^3$ | 88 | 30 |

Since *k*_on_ depends on the concentration of photoactivated molecules generated by pulse excitation, its absolute value is inherently ambiguous. As an example, we examined the sensitivity of the analysis to this parameter by performing additional calculations in which C₀ was varied from 0.3 µM to 10 µM (Fig. S6). The product *k*_on_·*C*₀ remained constant at 9 s^−1^, indicating a trade-off between the assumed concentration and the rate constant. This analysis suggests that the true value of *k*_on_ likely falls within the range of 0.9–30 µM^−1^s^−1^.

### 3.5 Reaction kinetics of wild-type OCP

Next, we analyzed the TG signals of wild-type (WT) OCP at various grating wavenumbers to investigate the contributions from both OCP^O^ monomer and dimer. As shown in Fig. 4(d), the signals exhibited complex *q*-dependent behaviors. At a high grating wavenumber (*q*^2^ = 7.3 × 10^12^ m^−2^), a monotonic decay was observed, indicating that the diffusion coefficient of early intermediates does not differ significantly from that of the reactant. In contrast, at an intermediate wavenumber (*q*^2^ = 6.1 × 10^11^ m^−2^), the thermal grating signal did not return to the baseline before a subsequent rise–decay feature appeared (Fig. 4(d), inset). The signs of refractive index changes of the signal suggest that the rising and decaying phases correspond to diffusion of the intermediate and the reactant, respectively. Such an increase in diffusion coefficient was not observed for OCP(R27L), suggesting that this phenomenon is unique to the presence of OCP^O^ dimers. While only a decrease in *D* due to dimerization was observed for the monomeric variant, the transient increase in *D* detected here may be attributed to the reverse reaction, that is, the dissociation of OCP^O^ dimers. A previous report by Rose *et al.* also proposed photo-induced dissociation of OCP^O^ dimers^49^; thus, the observed increase in diffusion coefficient could reasonably be interpreted as a result of OCP^O^ dimer dissociation.

At later times, signals attributable to the dimerization process become apparent. Given that the population of OCP^O^ dimers is high (approximately 80%) under the TG measurement conditions (100 µM), if the reaction were to terminate primarily with dimer dissociation, a strong dissociation-derived signal would be expected to persist. However, the pronounced association signal observed at later times indicates that the system does not simply end with dimer dissociation. Instead, it suggests that photoactivated species generated by dimer dissociation re-enter the association pathway and undergo dimerization once again.

Because the quantum yield of the OCP photoreaction is extremely low, it is highly unlikely that both monomers within a dimer are photoactivated by a single excitation pulse. Therefore, photoexcitation of the OCP^O^ dimer is expected to induce its dissociation into a photoactive monomer and an unreacted monomer. Based on these considerations, we constructed a reaction model in which the subsequent steps yield the OCP^R^ dimer as the final product, with the OCP^O^ monomer serving as a by-product. The reaction model was formalized as Scheme 2, where the subscript “d” signifies that the labeled species and associated rate constants originate from the photoactivation of the OCP^O^ dimer. The superscripts “M” and “D” denote whether each species is in the monomeric or dimeric state at the corresponding stage of the reaction scheme.

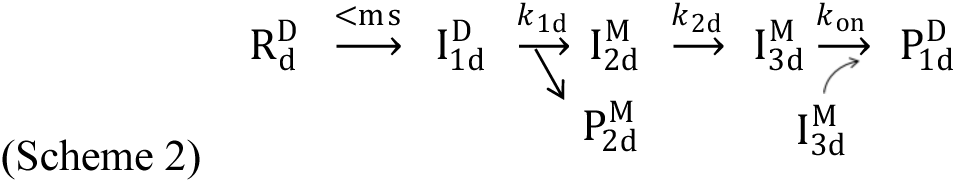

To analyze WT-OCP comprehensively, we constructed an integrated model (Scheme 3) that combines both the monomeric (Scheme 1) and dimeric (Scheme 2) pathways. In this scheme, R_m_ and R_d_ represent OCP^O^ monomer and dimer, respectively. Since the final product is the OCP^R^ dimer in both cases, we assumed that the intermediates formed after dissociation of the OCP^O^ dimer (I_2d_ and I_3d_) are identical to those formed in the reaction of the OCP^O^ monomer (I_2m_ and I_3m_). Because the signal attributed to dimer dissociation appeared at relatively early times, we further assumed that, following dissociation, the reaction pathway converges with that initiated from the OCP^O^ monomer. We also assumed that the photoexcited OCP molecules undergo the same sequence of reaction steps, so the reaction rate constants (*k*_2_ and *k*_on_) were set to be identical for both the monomer- and dimer-derived reactions.

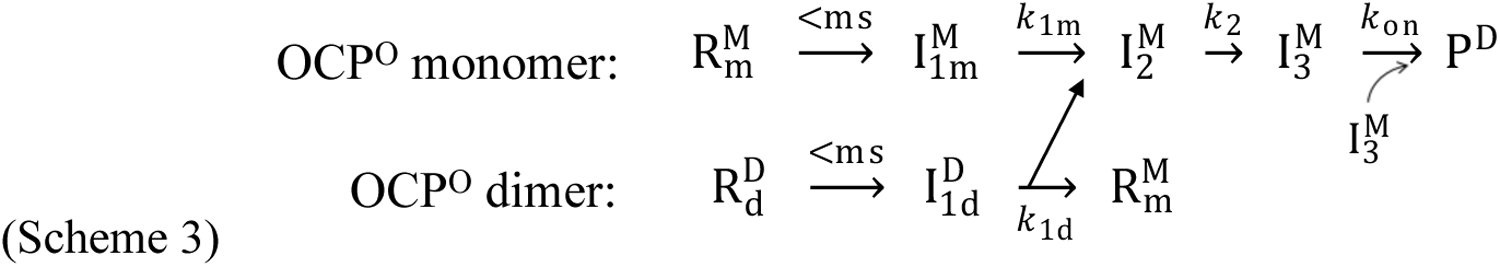

In the combined analysis, to reduce the arbitrariness of the fitting, the diffusion coefficient and kinetic parameters for the monomer reaction were fixed to the values obtained from OCP(R27L). The initial concentrations of OCP^O^ monomer and dimer were expressed using a fractional parameter. Numerical simulations were performed assuming a total photoactive concentration of *C*₀ = 0.3 μM. Despite the large number of fixed parameters, the model was able to successfully reproduce the observed signals, supporting the validity of the reaction model represented by Scheme 3. The newly obtained parameters for the reaction originating from the OCP^O^ dimer were also added to Table 2. The diffusion coefficient of the OCP^O^ dimer (*D*_R_ =7.66×10^−11^ m^2^s^−1^) was smaller than that of the monomer (*D*_R_ =8.53×10^−11^ m^2^s^−1^), consistent with the expected size difference. The reaction rate constant for the dissociation step was determined to be *k*₁_d_ = 1.0 × 10^3^ s^−1^. Notably, this time constant is comparable to that observed for the first structural transition in the monomeric mutant, suggesting that both pathways share the same rate-limiting step.

The fitted dimer fraction was 0.60, which is moderately lower than the theoretical estimate of 0.80 derived from SEC data. This discrepancy may indicate that the actual reactivity of OCP^O^ dimers is lower than that of OCP^O^ monomers, suggesting that the contribution of dimers to the observed photoreaction is smaller than would be expected based solely on their population.

### 3.6 Photo-reactivity of OCP^O^-monomer and OCP^O^-dimer

TG measurements suggest that dimerization in the dark reduces the photoreaction yield. To investigate the relationship between photo-reactivity and oligomeric state, we further analyzed the concentration dependence of light-induced absorbance changes under continuous illumination.

The accumulation of the light-adapted state (OCP^R^) was monitored by the change in absorbance at 550 nm immediately following continuous light irradiation (Fig. 5(a)(b)). The initial accumulation rate, was calculated from the change in absorbance (Δ*A*_550_) over the first 0.5 seconds using Eq. (5):

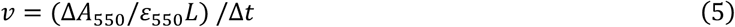

**Figure 5.**
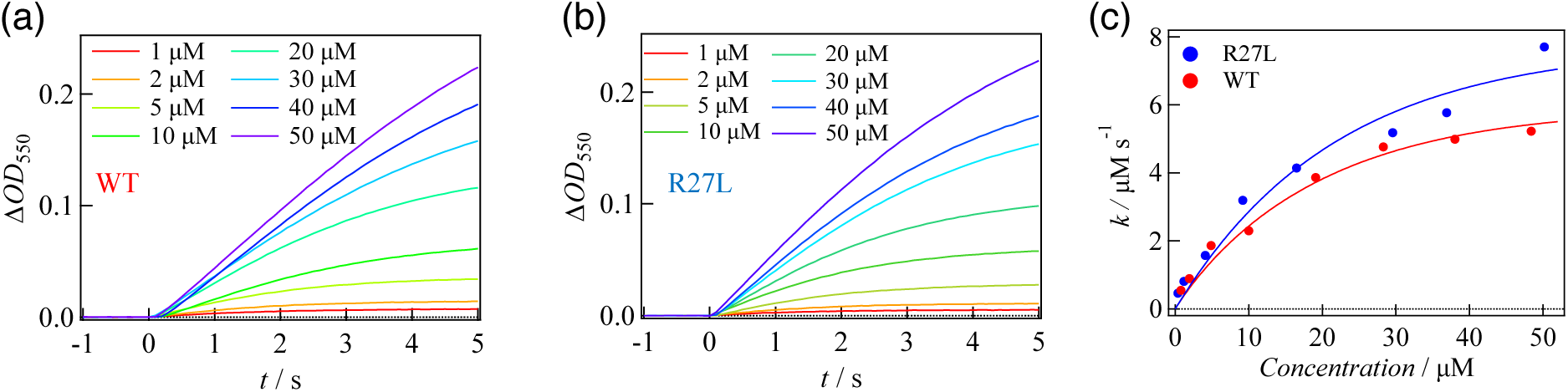
(a) Time-dependent change in absorbance at 550 nm for WT-OCP under continuous illumination with a blue LED. (b) Time-dependent change in absorbance at 550 nm for OCP(R27L) under the same conditions. Sample concentrations ranged from 1 μM to 50 μM, with illumination starting at t = 0 s. (c) Concentration dependence of the initial accumulation rates (initial velocities) for WT-OCP (red) and OCP(R27L) (blue). The solid lines represent the fitting curves using Eq. (S4.2) for WT and Eq. (S4.1) for the R27L mutant.

where *ε*_550_ is the extinction coefficient at 550 nm, Δ*t* is 0.5 s, and *L* is the optical path length for the absorbance measurement (1 mm).

First, we examined the concentration dependence of using the monomeric mutant OCP(R27L) (Fig. 5(c)). The data were fitted using Eq. (S4.1), yielding a rate constant for the monomer reaction of under the experimental conditions (7.8 *μ*M s^−1^).The same analysis was then performed for wild-type OCP^O^ at various concentrations. In WT-OCP samples, both monomeric and dimeric forms coexist and can contribute to OCP^R^ formation. At low concentrations (<5 µM), where the monomer is dominant, the initial accumulation rate was similar to that of OCP(R27L), confirming that the monomer reaction rate (7.8 *μ*M s^−1^) is similar between constructs.

The concentration dependence for WT was fitted using a two-component model incorporating contributions from both monomeric and dimeric species (Eq. (S4.2)). The fitting yielded rate constants of 7.8 *μ*M s^−1^ and 5.4 *μ*M s^−1^, respectively. The resulting ratio indicates that the photoreactivity of OCP^O^ dimers is approximately 30% lower than that of monomers. The consistently lower reactivity of the dimer species explains why its contribution to the TG signal was estimated to be smaller than expected from the population ratio alone, indicating that oligomerization reduces the quantum efficiency of the OCP photoreaction.

### 3.7 Heterodimer formation between apo-OCP and holo-OCP

In this section, we examine the effect of apo-OCP on the photoreaction behavior of OCP. First, the oligomeric state of apo-OCP was characterized using SEC and SAXS. As shown in Fig. S7(a), the SEC elution profile of apo-OCP gradually shifted toward larger molecular weights with increasing concentration, indicating concentration-dependent dimerization. From a plot of molecular mass versus concentration (Fig. S7(b)) and fitting using Eq. (3), the dissociation constant (*K*_d,apo_), monomer mass (mM), and dimer mass (dM) were determined to be 0.5 µM, 47 kDa, and 100 kDa, respectively. The low *K*_d,apo_ suggests that apo-OCP forms dimers much more stable than OCP^O^ or OCP^R^.

SAXS measurements further confirmed this interpretation. The scattering profile at 150 µM (Fig. S7(c)) closely resembled that of OCP^R^. Analysis using Guinier plots (Fig. S7(d)), *P*(r) distribution (Fig. S7(e)), and Porod volume calculations yielded *R*_g_ = 39.8 Å, *D*_max_ = 140 Å, and molecular mass ≈ 73 kDa. These values align with SEC results, indicating that apo-OCP predominantly exists as a dimer in solution. Structural modeling based on the scattering data (Fig. S7(f)) showed that the solution conformation of apo-OCP resembles that of the OCP^R^ dimer. This similarity is reasonable because apo-OCP lacks the carotenoid chromophore that links the NTD and CTD, which allows domain separation to occur. This is similar to what is observed in the light-adapted state of OCP.

Although the dimeric form of apo-OCP is stable, monomer–dimer equilibrium still exists. Therefore, apo-OCP monomers are always present in solution. Because apo-OCP monomers adopt an extended conformation similar to OCP^R^, they can potentially interact with OCP^R^ monomers to form heterodimers. To evaluate this possibility, we examined the TG signals of mixtures of OCP^O^ and apo-OCP.

In these experiments, the OCP^O^ concentration was fixed at 100 µM, while apo-OCP was added at concentrations ranging from 10 to 200 µM. Upon addition of 10 µM apo-OCP, the TG diffusion signal markedly increased. The signal intensity continued to rise with increasing apo-OCP concentration (Fig. 6(a)). Since apo-OCP lacks a chromophore and does not absorb light at the excitation wavelength, the enhancement in the diffusion signal must originate from the formation of heterodimers between OCP^R^ and apo-OCP. This result provides direct evidence that OCP^R^ monomers can associate with apo-OCP monomers (Fig. 6(b)).

**Figure 6.**
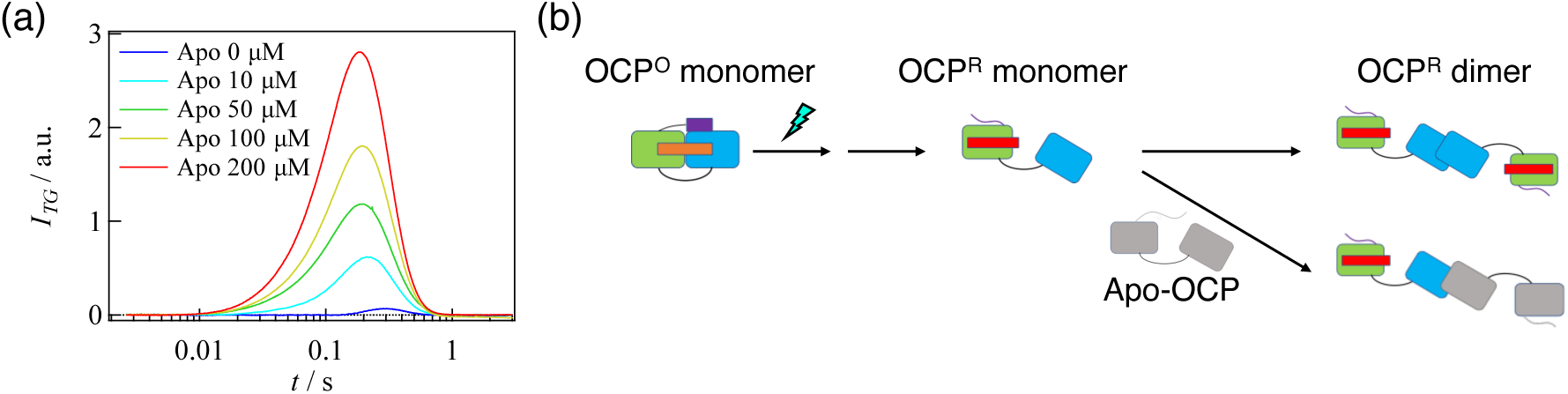
(a) TG signals of OCP (100 μM) measured in the presence of various concentrations of apo-OCP: 0 μM (purple), 10 μM (cyan), 50 μM (green), 100 μM (yellow), and 200 μM (red). The grating wavenumber was *q*^2^ = 1.2 × 10^11^ m^−2^. (b) Schematic illustration of the intermolecular interaction between the OCP^R^ monomer and apo-OCP. In the presence of apo-OCP, a side reaction involving heterodimer formation occurs.

## 4. Discussion

### 4.1 Reaction scheme of OCP

To elucidate the photo-induced dimerization mechanism of OCP, we integrate structural and kinetic findings to describe the transitions between reaction intermediates and the role of oligomeric state. SEC and SAXS measurements revealed that OCP exists in a monomer–dimer equilibrium under both dark- and light-adapted conditions, with OCP^R^ exhibiting a higher propensity for dimerization. Furthermore, SAXS data showed that OCP^R^ dimers adopt an elongated structure, while OCP^O^ dimers are more compact.

TG analysis of the monomeric mutant OCP(R27L) clarified the reaction pathway of the OCP^O^ monomer. Upon photoexcitation, a rapid initial reaction was followed by two additional intermediate steps before dimer formation, indicating that multiple sequential structural rearrangements precede the association of photoactivated species. The diffusion coefficient decreased in three distinct steps: the first occurred within 1 ms, the second followed at approximately 11 ms, and the final step, corresponding to OCP^R^ dimerization, occurred at around 100 ms with an estimated association rate constant of 30 μM^−1^s^−1^. The change in the diffusion coefficient was smallest in the first step, became somewhat larger in the second, and was most pronounced in the final step. These observations suggest that the initial step involves a local structural change, the second reflects a more substantial conformational rearrangement, and the final, largest change is associated with the dimerization reaction.

Previous studies^14,15,18^ have suggested that, after carotenoid translocation, the N-terminal extension (NTE) undergoes unfolding, followed by dissociation of the NTD and CTD domains. Based on this framework and the amplitudes of D-change at each step, we assign the 1 ms event to NTE dissociation and the 11 ms transition to the separation of NTD and CTD. These rearrangements likely expose interaction surfaces necessary for the subsequent dimerization step.

We then examined the behavior of OCP^O^ dimers. TG signal analysis allowed us to resolve the contribution of dimeric species, revealing that the OCP^O^ dimer dissociates upon photoexcitation into OCP^R^ and OCP^O^ monomers, which subsequently form OCP^R^ dimers. This result is consistent with the structural differences: while the OCP^O^ dimer adopts a compact conformation with an NTD–CTD interface, the OCP^R^ dimer associates via CTD–CTD interactions (Fig. 1(a)(b)). These differing interfaces imply that OCP^R^ dimers cannot form directly from the compact OCP^O^ dimer without structural rearrangement. Instead, the compact dimer must first dissociate into monomers that adopt an open conformation, enabling re-association through CTD–CTD contacts. This is further supported by the observation of OCP^O^-to-OCP^R^ dimer transition under continuous illumination in SAXS measurements, suggesting that this conversion proceeds via a monomeric intermediate.

Notably, the dissociation of the OCP^O^ dimer occurs with a time constant of ∼1 ms, coinciding with the NTE dissociation step observed in monomers. Given that the NTE is located at the dimer interface and forms a stabilizing salt bridge (D19–R27)^7^, its release likely destabilizes the dimer. This synchrony in kinetics between monomer and dimer pathways reinforces the interpretation that the first intermediate (I_2_) in the monomer reaction corresponds structurally to the NTE-unfolded state.

Importantly, the photo-reactivity of the OCP^O^ dimer is lower than that of the monomer, as revealed by both TG and continuous illumination experiments. This difference can be attributed to the restricted mobility of the NTE in the dimeric form, which is further stabilized by intermolecular interactions. Such structural constraints likely hinder the unfolding of NTE, which is essential for initiating downstream reactions.

Together, these findings suggest a comprehensive reaction scheme in which the OCP^O^ monomer undergoes an initial photoreaction followed by two sequential conformational changes prior to forming OCP^R^ dimers, while the OCP^O^ dimer must first dissociate to enter the same pathway. A schematic representation of the overall mechanism is shown in Fig. 7.

**Figure 7.**
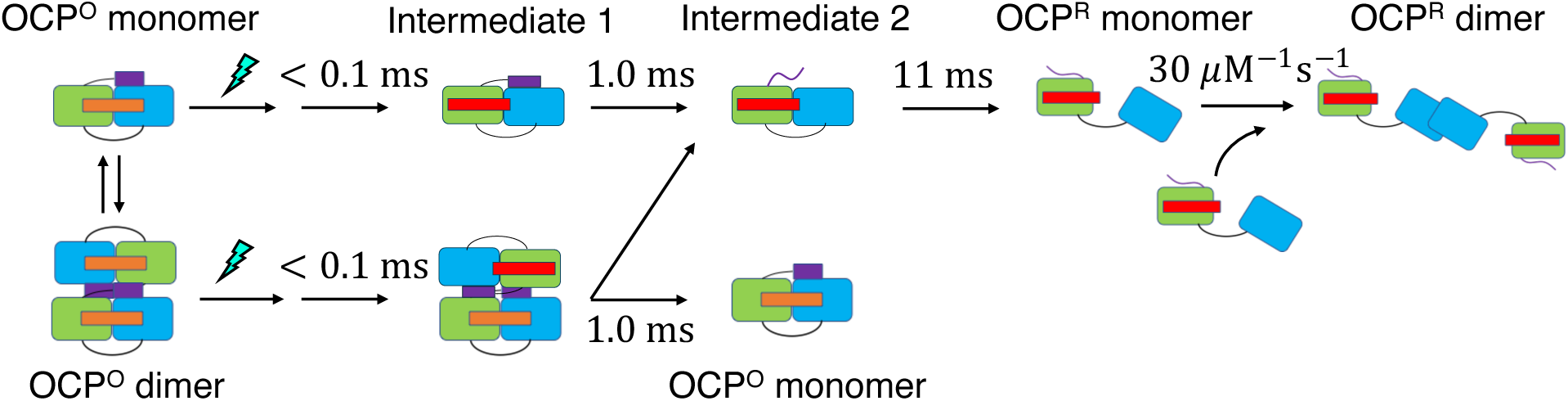
Proposed photoreaction scheme of OCP. In the dark state, OCP exists in equilibrium between monomeric and dimeric forms. Upon excitation of the monomer, a local photoreaction of the chromophore is followed by dissociation of the NTE and subsequent separation of the NTD and CTD domains, ultimately leading to dimerization of OCP^R^. When the dimeric form of OCP is excited, a local reaction of the chromophore also occurs, followed by dissociation into a photoactivated monomer and a dark-adapted monomer (OCP^O^ monomer). The photoactivated monomer then follows the same pathway as that initiated by light excitation of the OCP^O^ monomer.

### 4.2 Physiological significance of dimerization

OCP is a central protein in the non-photochemical quenching (NPQ) pathway of cyanobacteria, protecting the cell by dissipating excess energy as heat. However, excessive NPQ can reduce photosynthetic efficiency and limit growth. Therefore, properly controlling the activation threshold of OCP is important for optimizing photosynthetic performance under fluctuating light conditions.

Our results suggest that OCP^O^ dimer formation reduces the photo-reactivity of OCP. This attenuation may restrict OCP activation under low light conditions, ensuring that NPQ is triggered only under strong illumination. In other words, OCP^O^ dimerization may function as a molecular safeguard against excessive quenching. Conversely, the photoproduct OCP^R^ appears to function more effectively in its dimeric form. Cryo-EM structures of the OCP–PBS complex revealed that OCP^R^ dimers associate with both the core and rod regions of PBS. Furthermore, the dimer in the complex adopts an extended conformation, which closely matches the dimeric structure observed in solution in this study. This binding geometry suggests that dimerization enhances the structural stability of the OCP– PBS complex. These findings support the idea that OCP^R^ dimerization promotes more efficient NPQ activation upon photoactivation.

Taken together, these results indicate that dimerization in both dark- and light-adapted states may contribute to the fine-tuning of NPQ activity.

### 4.3 Influence of apo-OCP on data analyses

It has been found that apo-OCP readily forms dimers and can also form heterodimers with photoactivated holo-OCP^R^. This behavior significantly complicates the interpretation of experiments involving oligomerization and photoreaction analysis. Furthermore, because apo-OCP lacks the carotenoid chromophore, its incorporation into heterodimers is expected to reduce NPQ efficiency, potentially distorting the results of functional assays. Therefore, to ensure accurate and reproducible results in both mechanistic and functional studies, it is essential to completely remove apo-OCP from the sample prior to experimentation.

## 5. Conclusion

In this study, we investigated the photo-induced reaction dynamics of OCP and clarified in detail the kinetics of monomer- and dimer-initiated reactions. Following local photochemistry at the carotenoid site, the OCP^O^ monomer undergoes two sequential conformational transitions prior to forming OCP^R^ dimers. These steps likely correspond to displacement of the NTE and separation of the NTD and CTD domains, ultimately enabling dimerization of OCP^R^.

Interestingly, when the OCP^O^ dimer is photoexcited, it first dissociates into monomers before undergoing the same structural transitions, suggesting that the OCP^O^ and OCP^R^ dimers have distinct structural and interaction sites. Our results confirm this structural divergence and demonstrate that the dimeric form reduces the reaction yield, acting as a photoreactivity suppressor under low light conditions. Conversely, OCP^R^ homo-dimerization may enhance the stability of its interaction with PBS, highlighting a potential functional role in NPQ activation.

Given the growing interest in cyanobacteria as platforms for environmental and biotechnological applications, understanding the NPQ system, particularly the role of OCP, is of increasing importance. This study provides mechanistic insights into OCP function and lays the groundwork for future exploration of its interaction with regulatory proteins such as FRP and light-harvesting complexes such as phycobilisomes.

## Data availability

The data supporting this article have been included as part of the Supplementary Information.

## Acknowledgements

The authors are grateful for the generous funding support provided to M.T. (JSPS KAKENHI 19H01863, 21H01885, 21K19218) and to Y.N. (JSPS KAKENHI JP22K18361; Toyota Riken Scholar; Shimadzu Science Foundation; SPIRIT2 of Kyoto University).

## References

(1) Triantaphylidès, C.; Havaux, M. Singlet Oxygen in Plants: Production, Detoxification and Signaling. Trends in Plant Science. April 2009, pp 219–228. 10.1016/j.tplants.2009.01.008.

(2) Magdaong, N. C. M.; Blankenship, R. E. Photoprotective, Excited-State Quenching Mechanisms in Diverse Photosynthetic Organisms. Journal of Biological Chemistry. American Society for Biochemistry and Molecular Biology Inc. April 6, 2018, pp 5018–5025. 10.1074/jbc.TM117.000233.

(3) Goss, R.; Lepetit, B. Biodiversity of NPQ. Journal of Plant Physiology. Urban und Fischer Verlag GmbH und Co. KG January 1, 2015, pp 13–32. 10.1016/j.jplph.2014.03.004.

(4) Kirilovsky, D. Photoprotection in Cyanobacteria: The Orange Carotenoid Protein (OCP)-Related Non-Photochemical-Quenching Mechanism. Photosynthesis Research. July 2007, pp 7–16. 10.1007/s11120-007-9168-y.

(5) Sluchanko, N. N.; Slonimskiy, Y. B.; Maksimov, E. G. Features of Protein −protein Interactions in the Cyanobacterial Photoprotection Mechanism. Biochemistry (Moscow). December 1, 2017, pp 1592–1614. 10.1134/s000629791713003x.

(6) Holt, T. K.; Krogmann, D. W. A CAROTENOID-PROTEIN FROM CYANOBACTERIA; 1981; Vol. 637, pp 408–414.

(7) Wilson, A.; Kinney, J. N.; Zwart, P. H.; Punginelli, C.; D’Haene, S.; Perreau, F.; Klein, M. G.; Kirilovsky, D.; Kerfeld, C. A. Structural Determinants Underlying Photoprotection in the Photoactive Orange Carotenoid Protein of Cyanobacteria. J. Biol. Chem. 2010, 285 (24), 18364–18375.

(8) Chukhutsina, V. U.; Baxter, J. M.; Fadini, A.; Morgan, R. M.; Pope, M. A.; Maghlaoui, K.; Orr, C. M.; Wagner, A.; van Thor, J. J. Light Activation of Orange Carotenoid Protein Reveals Bicycle-Pedal Single-Bond Isomerization. Nat. Commun. 2022, 13 (1). 10.1038/s41467-022-34137-4.

(9) Fujisawa, T.; Leverenz, R. L.; Nagamine, M.; Kerfeld, C. A.; Unno, M. Raman Optical Activity Reveals Carotenoid Photoactivation Events in the Orange Carotenoid Protein in Solution. J. Am. Chem. Soc. 2017, 139 (30), 10456–10460.

(10) Wilson, A.; Punginelli, C.; Couturier, M.; Perreau, F.; Kirilovsky, D. Essential Role of Two Tyrosines and Two Tryptophans on the Photoprotection Activity of the Orange Carotenoid Protein. Biochim. Biophys. Acta Bioenerg. 2011, 1807 (3), 293–301.

(11) Leverenz, R. L.; Sutter, M.; Wilson, A.; Gupta, S.; Thurotte, A.; Bourcier de Carbon, C.; Petzold, C. J.; Ralston, C.; Perreau, F.; Kirilovsky, D.; Kerfeld, C. A. A 12 Å Carotenoid Translocation in a Photoswitch Associated with Cyanobacterial Photoprotection. Science 2015, 348 (6242), 1463–1466.

(12) Pigni, N. B.; Clark, K. L.; Beck, W. F.; Gascon, J. A. Spectral Signatures of Canthaxanthin Translocation in the Orange Carotenoid Protein. J. Phys. Chem. B 2020, 124 (50), 11387–11395.

(13) Niziński, S. N.; Wilson, A.; Uriarte, L. M.; Ruckebusch, C.; Andreeva, E. A.; Schlichting, I.; Colletier, J.-P.; Kirilovsky, D.; Burdzinski, G.; Sliwa, M. Unifying Perspective of the Ultrafast Photodynamics of Orange Carotenoid Proteins from Synechocystis: Peril of High-Power Excitation, Existence of Different S* States, and Influence of Tagging. J. Am. Chem. Soc. 2022, 2, 1084–1095.

(14) Sharawy, M.; Pigni, N. B.; May, E. R.; Gascón, J. A. A Favorable Path to Domain Separation in the Orange Carotenoid Protein. Protein Sci. 2022, 31 (4), 850–863.

(15) Konold, P. E.; Van Stokkum, I. H. M.; Muzzopappa, F.; Wilson, A.; Groot, M. L.; Kirilovsky, D.; Kennis, J. T. M. Photoactivation Mechanism, Timing of Protein Secondary Structure Dynamics and Carotenoid Translocation in the Orange Carotenoid Protein. J. Am. Chem. Soc. 2019, 141 (1), 520–530.

(16) Sluchanko, N. N.; Klementiev, K. E.; Shirshin, E. A.; Tsoraev, G. V.; Friedrich, T.; Maksimov, E. G. The Purple Trp288Ala Mutant of Synechocystis OCP Persistently Quenches Phycobilisome Fluorescence and Tightly Interacts with FRP. Biochim. Biophys. Acta Bioenerg. 2017, 1858 (1), 1–11.

(17) Maksimov, E. G.; Sluchanko, N. N.; Slonimskiy, Y. B.; Slutskaya, E. A.; Stepanov, A. V.; Argentova-Stevens, A. M.; Shirshin, E. A.; Tsoraev, G. V.; Klementiev, K. E.; Slatinskaya, O. V.; Lukashev, E. P.; Friedrich, T.; Paschenko, V. Z.; Rubin, A. B. The Photocycle of Orange Carotenoid Protein Conceals Distinct Intermediates and Asynchronous Changes in the Carotenoid and Protein Components. Sci. Rep. 2017, 7 (1). 10.1038/s41598-017-15520-4.

(18) Mezzetti, A.; Alexandre, M.; Thurotte, A.; Wilson, A.; Gwizdala, M.; Kirilovsky, D. Two-Step Structural Changes in Orange Carotenoid Protein Photoactivation Revealed by Time-Resolved Fourier Transform Infrared Spectroscopy. J. Phys. Chem. B 2019, 123 (15), 3259–3266.

(19) Maksimov, E. G.; Protasova, E. A.; Tsoraev, G. V.; Yaroshevich, I. A.; Maydykovskiy, A. I.; Shirshin, E. A.; Gostev, T. S.; Jelzow, A.; Moldenhauer, M.; Slonimskiy, Y. B.; Sluchanko, N. N.; Friedrich, T. Probing of Carotenoid-Tryptophan Hydrogen Bonding Dynamics in the Single-Tryptophan Photoactive Orange Carotenoid Protein. Sci. Rep. 2020, 10 (1). 10.1038/s41598-020-68463-8.

(20) Liu, H.; Zhang, H.; King, J. D.; Wolf, N. R.; Prado, M.; Gross, M. L.; Blankenship, R. E. Mass Spectrometry Footprinting Reveals the Structural Rearrangements of Cyanobacterial Orange Carotenoid Protein upon Light Activation. Biochim. Biophys. Acta Bioenerg. 2014, 1837 (12), 1955–1963.

(21) Liu, H.; Zhang, H.; Orf, G. S.; Lu, Y.; Jiang, J.; King, J. D.; Wolf, N. R.; Gross, M. L.; Blankenship, R. E. Dramatic Domain Rearrangements of the Cyanobacterial Orange Carotenoid Protein upon Photoactivation. Biochemistry 2016, 55 (7), 1003–1009.

(22) Gupta, S.; Guttman, M.; Leverenz, R. L.; Zhumadilova, K.; Pawlowski, E. G.; Petzold, C. J.; Lee, K. K.; Ralston, C. Y.; Kerfeld, C. A. Local and Global Structural Drivers for the Photoactivation of the Orange Carotenoid Protein. Proc. Natl. Acad. Sci. U. S. A. 2015, 112 (41), E5567–E5574.

(23) Andreeva, E. A.; Niziński, S.; Wilson, A.; Levantino, M.; De Zitter, E.; Munro, R.; Muzzopappa, F.; Thureau, A.; Zala, N.; Burdzinski, G.; Sliwa, M.; Kirilovsky, D.; Schirò, G.; Colletier, J. P. Oligomerization Processes Limit Photoactivation and Recovery of the Orange Carotenoid Protein. Biophys. J. 2022, 121 (15), 2849–2872.

(24) Golub, M.; Moldenhauer, M.; Schmitt, F. J.; Feoktystov, A.; Mändar, H.; Maksimov, E.; Friedrich, T.; Pieper, J. Solution Structure and Conformational Flexibility in the Active State of the Orange Carotenoid Protein: Part I. Small-Angle Scattering. J. Phys. Chem. B 2019, 123 (45), 9525–9535.

(25) Golub, M.; Moldenhauer, M.; Matsarskaia, O.; Martel, A.; Grudinin, S.; Soloviov, D.; Kuklin, A.; Maksimov, E. G.; Friedrich, T.; Pieper, J. Stages of OCP-FRP Interactions in the Regulation of Photoprotection in Cyanobacteria, Part 2: Small-Angle Neutron Scattering with Partial Deuteration. J. Phys. Chem. B 2023, 127 (9), 1901–1913.

(26) Domínguez-Martín, M. A.; Sauer, P. V.; Kirst, H.; Sutter, M.; Bína, D.; Greber, B. J.; Nogales, E.; Polívka, T.; Kerfeld, C. A. Structures of a Phycobilisome in Light-Harvesting and Photoprotected States. Nature 2022, 609 (7928), 835–845.

(27) Moldenhauer, M.; Sluchanko, N. N.; Tavraz, N. N.; Junghans, C.; Buhrke, D.; Willoweit, M.; Chiappisi, L.; Schmitt, F. J.; Vukojević, V.; Shirshin, E. A.; Ponomarev, V. Y.; Paschenko, V. Z.; Gradzielski, M.; Maksimov, E. G.; Friedrich, T. Interaction of the Signaling State Analog and the Apoprotein Form of the Orange Carotenoid Protein with the Fluorescence Recovery Protein. Photosynth. Res. 2018, 135 (1–3), 125–139.

(28) Nakasone, Y.; Kawasaki, Y.; Konno, M.; Inoue, K.; Terazima, M. Time-Resolved Detection of Light-Induced Conformational Changes of Heliorhodopsin. Phys. Chem. Chem. Phys. 2023, 25 (18), 12833–12840.

(29) Terazima, M. Studies of Photo-Induced Protein Reactions by Spectrally Silent Reaction Dynamics Detection Methods: Applications to the Photoreaction of the LOV2 Domain of Phototropin from Arabidopsis Thaliana. Biochimica et Biophysica Acta - Proteins and Proteomics. August 2011, pp 1093–1105. 10.1016/j.bbapap.2010.12.011.

(30) Terazima, M. Spectrally Silent Protein Reaction Dynamics Revealed by Time-Resolved Thermodynamics and Diffusion Techniques. Acc. Chem. Res. 2021, 54 (9), 2238–2248.

(31) Terazima, M. Time-Resolved Detection of Association/Dissociation Reactions and Conformation Changes in Photosensor Proteins for Application in Optogenetics. Biophysical Reviews. Springer Science and Business Media Deutschland GmbH December 1, 2021, pp 1053–1059. 10.1007/s12551-021-00868-9.

(32) Terazima, M. Time-Dependent Intermolecular Interaction during Protein Reactions. Physical Chemistry Chemical Physics. October 14, 2011, pp 16928–16940. 10.1039/c1cp21868a.

(33) Muzzopappa, F.; Wilson, A.; Kirilovsky, D. Interdomain Interactions Reveal the Molecular Evolution of the Orange Carotenoid Protein. Nat. Plants 2019, 5 (10), 1076–1086.

(34) Li, X. D.; Zhou, L. J.; Zhao, C.; Lu, L.; Niu, N. N.; Han, J. X.; Zhao, K. H. Optimization of Expression of Orange Carotenoid Protein in Escherichia Coli. Protein Expr. Purif. 2019, 156, 66‒71.

(35) Chen, Y. W. Structural Genomics; Vol. 1091. http://www.springer.com/series/7651.

(36) Manalastas-Cantos, K.; Konarev, P. V.; Hajizadeh, N. R.; Kikhney, A. G.; Petoukhov, M. V.; Molodenskiy, D. S.; Franke. ATSAS 3.0: Expanded Functionality and New Tools for Small-Angle Scattering Data Analysis. Journal of applied crystallography 2021, 54 (1), 343–355.

(37) Franke, D.; Petoukhov, M. V.; Konarev, P. V.; Panjkovich, A.; Tuukkanen, A.; Mertens, H. D. T.; Kikhney, A. G.; Hajizadeh, N. R.; Franklin, J. M.; Jeffries, C. M.; Svergun, D. I. ATSAS 2.8: A Comprehensive Data Analysis Suite for Small-Angle Scattering from Macromolecular Solutions. J. Appl. Crystallogr. 2017, 50 (Pt 4), 1212–1225.

(38) Svergun, D. I. Determination of the Regularization Parameter in Indirect-Transform Methods Using Perceptual Criteria. Journal of applied crystallography 1992, 25 (4), 495–503.

(39) Svergun, D. I.; Petoukhov, M. V.; Koch, M. H. J. Determination of Domain Structure of Proteins from X-Ray Solution Scattering. Biophys. J. 2001, 80 (6), 2946–2953.

(40) Porod, G. General Theory. In Small angle X-ray scattering; inis.iaea.org, 1982.

(41) Brookes, E.; Rocco, M. Recent Advances in the UltraScan SOlution MOdeller (US-SOMO) Hydrodynamic and Small-Angle Scattering Data Analysis and Simulation Suite. Eur. Biophys. J. 2018, 47 (7), 855–864.

(42) Kim, S.; Nakasone, Y.; Takakado, A.; Yamazaki, Y.; Kamikubo, H.; Terazima, M. A Unique Photochromic UV-A Sensor Protein, Rc-PYP, Interacting with the PYP-Binding Protein. Phys. Chem. Chem. Phys. 2021, 23 (33), 17813–17825.

(43) Takekawa, G.; Nakasone, Y.; Kamiya, Y.; Asanuma, H.; Terazima, M. Reaction and Interaction Dynamics of Azobenzene-Tethered DNA with T7 RNA Polymerase. Phys. Chem. Chem. Phys. 2025, 27 (6), 3302–3312.

(44) Nakasone, Y.; Murakami, H.; Tokonami, S.; Oda, T.; Terazima, M. Time-Resolved Study on Signaling Pathway of Photoactivated Adenylate Cyclase and Its Nonlinear Optical Response. J. Biol. Chem. 2023, 299 (11), 105285.

(45) Shibata, K.; Nakasone, Y.; Terazima, M. Selective Photoinduced Dimerization and Slow Recovery of a BLUF Domain of EB1. J. Phys. Chem. B 2022, 126 (5), 1024–1033.

(46) Choi, S.; Nakasone, Y.; Hellingwerf, K. J.; Terazima, M. Photoreaction Dynamics of a Full-Length Protein YtvA and Intermolecular Interaction with RsbRA. Biochemistry 2020, 59 (50), 4703–4710.

(47) Iwata, K.; Terazima, M.; Masuhara, H. Novel Physical Chemistry Approaches in Biophysical Researches with Advanced Application of Lasers: Detection and Manipulation. Biochimica et Biophysica Acta - General Subjects. Elsevier B.V. February 1, 2018, pp 335–357. 10.1016/j.bbagen.2017.11.003.

(48) Nemoto, N.; Koike, A.; Osaki, K.; Koseki, T.; Doi, E. Dynamic Light Scattering of Aqueous Solutions of Linear Aggregates Induced by Thermal Denaturation of Ovalbumin. Biopolymers 1993, 33 (4), 551–559.

(49) Rose, J. B.; Gascón, J. A.; Sutter, M.; Sheppard, D. I.; Kerfeld, C. A.; Beck, W. F. Photoactivation of the Orange Carotenoid Protein Requires Two Light-Driven Reactions Mediated by a Metastable Monomeric Intermediate. Phys. Chem. Chem. Phys. 2023, 25 (48), 33000–33012.

