## Supplementary information for "Kinetic Insights into Photoinduced Monomer–Dimer Conversion and Activation of Orange Carotenoid Protein"

### Supporting Information

Tadayuki Tokashiki[1], Takatoshi Ohata[1], Shunrou Tokonami[2], Takashi Oda[3], Yusuke Nakasone[1], Masahide Terazima[1]

[1] Department of Chemistry, Graduate School of Science, Kyoto University, Kyoto, Japan

[2] Department of Chemistry, Faculty of Science, Gakushuin University, Tokyo, Japan

[3] J-PARC Center, JAEA, Ibaraki, Japan

#### S1. SEC profiles

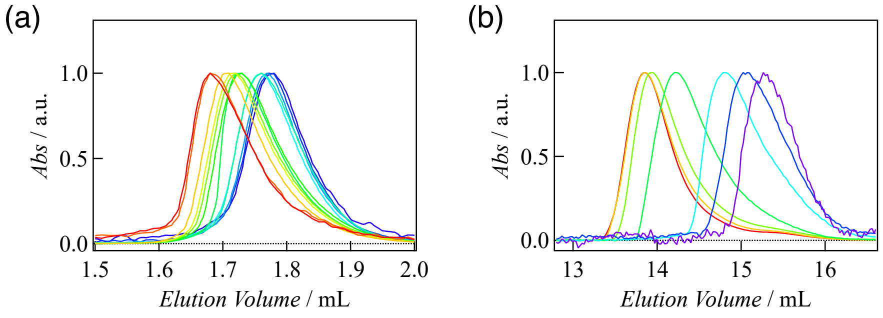


Figure S1. SEC profiles of OCP measured at 280 nm for various sample concentrations, with each elution profile normalized to its peak intensity. (a) SEC profiles in the dark-adapted state. Sample concentrations ranged from 400 μM (red) to 0.5 μM (purple). (b) SEC profiles under illumination. Sample concentrations ranged from 300 μM (red) to 1 μM (purple).

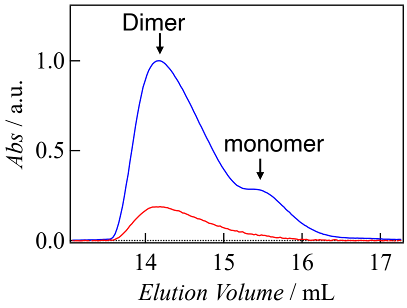


Figure S2. SEC profiles under conditions where $OCP^O$ and $OCP^R$ coexist. The intensity of the Xe lamp used for illumination was reduced to approximately 5% of that used in the other light-adapted measurements. The red and blue chromatograms represent monitoring at 620 nm and 280 nm, respectively.

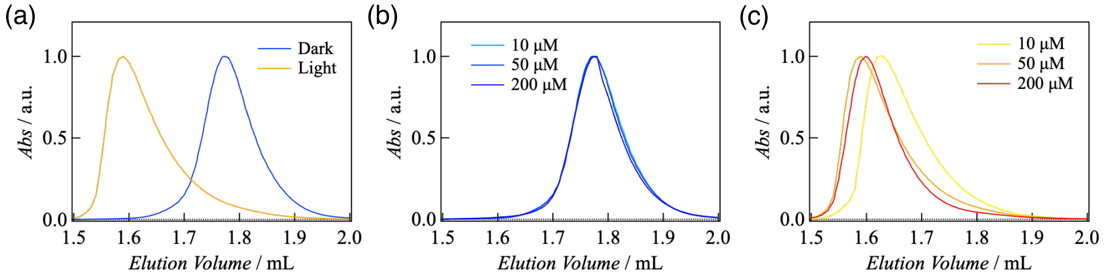


Figure S3. SEC profiles of the R27L mutant, normalized to peak intensity. (a) Comparison between dark- and light-adapted states at an injection concentration of 50 μM. (b) Concentration dependence in the dark-adapted state. (c) Concentration dependence in the light-adapted state.

**S2. SAXS analysis**

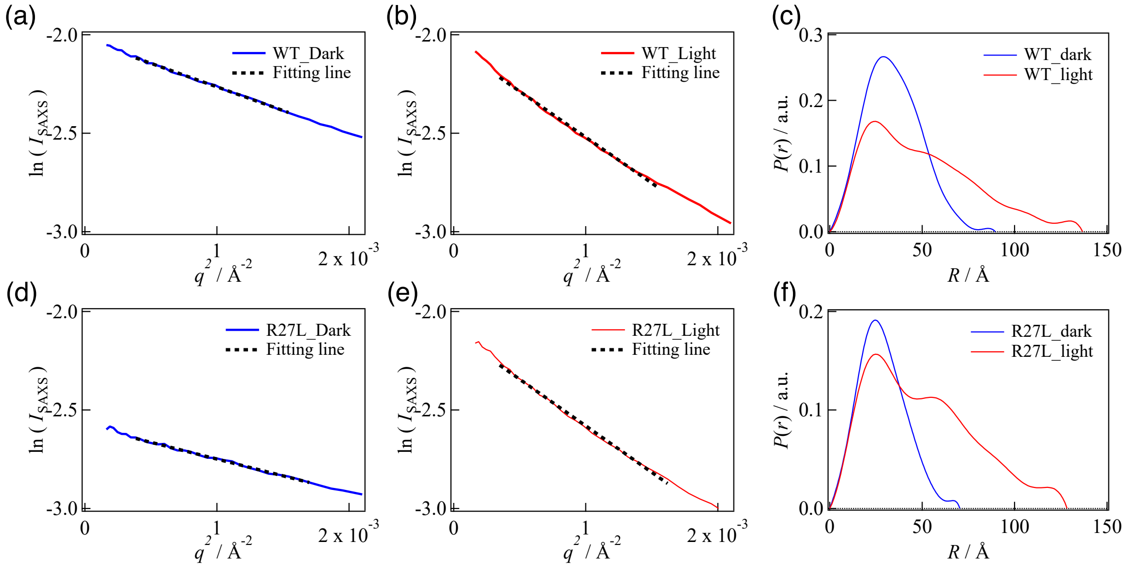


Figure S4. (a), (b) Guinier analysis of SAXS profiles for WT-OCP. The dotted line represents the fitting line. (c) Radial distribution function of WT-OCP. (d), (e) Guinier analysis of SAXS profiles for the R27L mutant. The dotted line represents the fitting line. (f) Radial distribution function of the R27L mutant.

### S3. TG signals by successive pulse-excitations

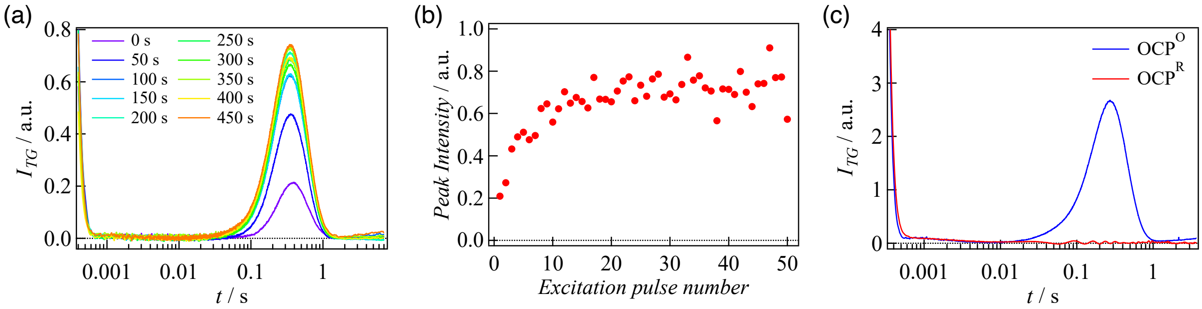


Figure S5. (a) Temporal evolution of TG signals under repeated pulse excitation at 0.1 Hz. Seven independent excitation experiments were performed, and the corresponding pulse-induced TG signals were averaged. The grating wavenumber was $q^2 = 5.5 \times 10^{10}$ m$^{-2}$. (b) Peak intensity of the diffusion signals shown in (a). (c) Comparison of TG signals under conditions where $OCP^R$ (red) or $OCP^O$ (blue) is dominant. These results indicate that $OCP^R$ does not contribute to the diffusion signal.

**S4. Accumulation rates under continuous light illumination**

Initial accumulation rate of $OCP^R$ was measured after starting continuous light illumination using a blue LED. The LED light was irradiated onto the cell at an incident angle of 60°. The sample cell path length was set to 1 mm, ensuring that the absorbance at 460 nm from blue LED light remained approximately 0.6 even at a concentration of 50 $\mu$M. The absorbance change at $\lambda$=550 nm was monitored, and the accumulation rate, $v$, was calculated using Eq. (4) as the initial slope of the absorbance change.

First, OCP(R27L), which exists predominantly as a monomer was used to determine the rate constant of $OCP^O$ monomer. The initial accumulation rate should be proportional to the absorbed light intensity. According to the Lambert-Beer's law, the absorbed light intensity ($I_{abs}$) is given by

$$I_{\rm abs} = I_0(1 - 10^{-\varepsilon C L_{ex}})$$

where $I_0$, $\varepsilon$, $C$ and $L_{ex}$ represent as the incident light intensity, molar extinction coefficient, concentration, and path length for excitation, respectively. Hence, the accumulation rate of the product ($v$) may be given by Eq. (S4.1);

$$v = v_{\rm m}\,(1 - 10^{-\varepsilon C L_{ex}}) \qquad \text{(S4.1)}$$

where $v_m$ represents the accumulation rate of the monomer under the experimental excitation conditions. Using this equation, the observed accumulation rate of OCP(R27L) at various concentrations (Fig. 5(c)) was fitted using the extinction coefficient at 460 nm $\varepsilon_{460}$ = 0.1 $\mu M^{-1}$ $cm^{-1}$ calculated using the extinction coefficient at 476 nm $\varepsilon_{476}$ = 0.118 $\mu M^{-1}$ $cm^{-1}$ and the absorbance ratio between 460 nm and 476 nm in the dark adapted state (Fig.1(c)), and $L_{ex}$ = 2 mm, which is calculated from the incident angle ($\theta$=60°) and the sample cell thickness of 1 mm, and an adjustable parameter $v_m$. The best fit was obtained for $v_m$= 7.76 $\mu$M $s^{-1}$.

For WT-OCP, there are two reactants, $OCP^O$-monomer and $OCP^O$-dimer, and the accumulation rates may be given by Eq. (S4.2).

$$v = \left(v_{\rm m}\frac{[{\rm OCP_m^O}]}{C} + v_{\rm d}\frac{2[{\rm OCP_d^O}]}{C}\right)(1 - 10^{-\varepsilon C L}) \qquad \text{(S4.2)}$$

where $v_d$ represents the accumulation rate of $OCP^O$-dimer. Furthermore, $[{\rm OCP_m^O}]$ and $[{\rm OCP_d^O}]$ are the concentrations of $OCP^O$-monomer and dimer, respectively. Using the same experimental conditions as that of OCP(R27L), the extinction coefficient $\varepsilon$ and the optical path length $L$ should be the same as those for OCP(R27L). $[{\rm OCP_m^O}]$ and $[{\rm OCP_d^O}]$ are calculated by Eq. (S1.3) and the dissociation constant of $OCP^O$-dimer, 10 $\mu$M was used. The best fit was obtained for $v_d$= 5.44 $\mu$M $s^{-1}$ when $v_m$ was fixed to be $v_m$= 7.76 $\mu$M $s^{-1}$.

**S5. Numerical calculation of TG signals for OCP(R27L)**

Based on the reaction scheme1, the reaction-diffusion equations can be expressed as Eq. (S5.1) – (S5.5). These differential equations cannot be solved analytically. To determine the kinetic parameters including the association rate constant between the photoexcited OCP molecules, numerical calculations were conducted to reproduce the TG signals. The calculation methods were shown below.

First, diffusion-reaction equations based on the reaction model (Scheme 1) can be depicted as following equations, where $C_X$ (x,t) represents the concentration of a species X at the position of $x$ and the time of $t$.

$$\frac{\partial C_R(x,t)}{\partial t} = D_R \frac{\partial^2 C_R(x,t)}{\partial x^2} \tag{S5.1}$$

$$\frac{\partial C_{I_1}(x,t)}{\partial t} = D_{I_1} \frac{\partial^2 C_{I_1}(x,t)}{\partial x^2} - k_1 C_{I_1}(x,t) \tag{S5.2}$$

$$\frac{\partial C_{I_2}(x,t)}{\partial t} = D_{I_2} \frac{\partial^2 C_{I_2}(x,t)}{\partial x^2} + k_1 C_{I_1}(x,t) - k_2 C_{I_2}(x,t) \tag{S5.3}$$

$$\frac{\partial C_{I_3}(x,t)}{\partial t} = D_{I_3} \frac{\partial^2 C_{I_3}(x,t)}{\partial x^2} + k_2 C_{I_2}(x,t) - 2k_{on} C_{I_3}^2(x,t) \tag{S5.4}$$

$$\frac{\partial C_P(x,t)}{\partial t} = D_P \frac{\partial^2 C_P(x,t)}{\partial x^2} + k_{on} C_{I_3}^2(x,t) \tag{S5.5}$$

When the crossed pump pulses excited the sample solution sinusoidally, the initial perturbation of the concentration of each species can be represent as Eq. (S5.6) to (S5.10).

$$\Delta C_R(x,0) = -C_0(1 - \cos(qx)) \tag{S5.6}$$

$$\Delta C_{I_1}(x,0) = C_0(1 - \cos(qx)) \tag{S5.7}$$

$$\Delta C_{I_2}(x,0) = 0 \tag{S5.8}$$

$$\Delta C_{I_3}(x,0) = 0 \tag{S5.9}$$

$$\Delta C_P(x,0) = 0 \tag{S5.10}$$

For general TG analyses, the analytical equation for TG signals can be acquired by solving the diffusion-reaction equations with the initial conditions using Fourie transforms and Laplace transforms. For these diffusion-reaction equations, however, we cannot solve them analytically. Then, numerical calculations are needed to acquire the time-evolutions of the concentration of each species.

The terms in the second derivative of the right-hand side of the reaction-diffusion equation can be approximated as Eq. (S5.13) using the central-difference approximation shown below.

the second derivative terms can be approximated as Eq. (S5.11).

$$\frac{d^2 f(x)}{dx^2} \approx \frac{f(x+h) + f(x-h) - 2f(x)}{h^2} \tag{S5.11}$$

$\partial C_X(x,t)/\partial t$ was calculated at each time t using this approximation, and the increment of the concentration was acquired by multiplying a time increment $\Delta t$. Repeating this process enables to obtain the time-evolutions of the concentration $C_X$ $(x,t)$.

Then, Fourie transformation was performed on $C_X$ $(x,t)$ to obtain the time evolution of concentration modulation $\widetilde{C_X}(\nu, t)$ corresponding to each frequency component.

$$\widetilde{C_X}(\nu, t) = \mathcal{F}[C_X(x, t)] \tag{S5.12}$$

To note, the only components that contribute to the formation of the TG signal are those that satisfy the phase matching condition with the probe light, which are the frequency components satisfying $\nu = q/2\pi$. Therefore, we extracted the satisfying component $\widetilde{C_X}(\nu = q/2\pi, t)$ from the obtained $\widetilde{C_X}(\nu, t)$ and used it as the TG generative component. When the diffractive index of each species X is expressed as $\delta n^0{}_X$, the total diffractive index $\delta n(t)$ can be depicted as:

$$\delta n(t) = \delta n_R^0 \widetilde{C_R}(t) + \delta n_{I_1}^0 \widetilde{C_{I_1}}(t) + \delta n_{I_2}^0 \widetilde{C_{I_2}}(t) + \delta n_P^0 \widetilde{C_P}(t) \tag{S5.13}$$

Since the signal intensity of the TG signal is proportional to the squared diffractive index, the time evolution of the signal intensity can be expressed as Eq. (S5.14)

$$I_{TG} = \alpha\left\{\delta n_R^0 \widetilde{C_R}(t) + \delta n_{I_1}^0 \widetilde{C_{I_1}}(t) + \delta n_{I_2}^0 \widetilde{C_{I_2}}(t) + \delta n_P^0 \widetilde{C_P}(t)\right\}^2 \tag{S5.14}$$

From the above calculations, the numerical calculations for the time evolution of the TG signals based on the reaction model including an association reaction between excited molecules were successfully performed. For this calculation, we need to input the initial value of the concentrations $C_0$ and the parameters of the proportional constant of $\alpha$, the grating wavenumber of $q$ and the diffusion coefficients of $D_X$ and the diffractive indexes of the transient species X.

In this calculation, the concentration of the excited molecules was set to $C_0$=0.3 $\mu$M, which was calculated by multiplying the total concentration ($C$=100 $\mu$M) and the reaction quantum yield of the photoreaction ($\Phi$=0.003). Numerical simulations were performed by varying the parameters to reproduce the TG signals of OCP(R27L) at the different grating wavenumbers, and the rate constants and diffusion coefficients were obtained (Table 2).

In the present calculation, the initial concentration $C_0$ was estimated under the assumption that each OCP molecule undergoes the photoreaction only once per excitation pulse. However, since the excitation pulse has a duration on the order of nanoseconds, it is possible that molecules which relax without reacting may be re-excited within the same pulse. Therefore, the actual initial concentration $C_0$ is likely somewhat higher than the estimated value. To examine the effect of initial concentration on the association rate constant $k_{on}$ obtained from numerical calculations, the analysis was repeated using different initial concentrations. The best agreement with the experimental signal was observed when $k_{on}C_0$=9 $s^{-1}$, indicating that this condition most accurately reproduced the measured TG signal (Fig. S7(b)). Therefore, since the actual concentration of reactive molecules was likely larger than 0.3 μM, the accurate value of the association rate should be slightly smaller than 30 $\mu M^{-1} s^{-1}$, but this analysis provided a reasonable estimate of its magnitude.

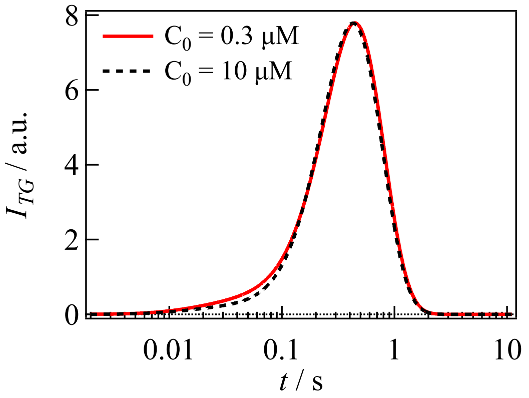


Figure S6. Calculated signals using Scheme 1 with different initial concentrations of photoactivated molecules ($C_0$). The red curve was obtained using $C_0$ = 0.3 μM, and the dotted curve using $C_0$ = 10 μM. The association rate constant was $k_{on}$ = 30 μM$^{-1}$s$^{-1}$ for $C_0$ = 0.3 μM and $k_{on}$ = 0.9 μM$^{-1}$s$^{-1}$ for $C_0$ = 10 μM; that is, the product of $k_{on}$ and $C_0$ remained constant at 9 s$^{-1}$.

**S6. Numerical calculation of TG signals for WT-OCP**

The numerical calculation for WT-OCP were conducted in the same way as for OCP(R27L). Based on Scheme 3, the reaction-diffusion equations can be expressed as Eq. (S6.1) – (S6.7).

$$\frac{\partial C_{R_1}(x,t)}{\partial t} = D_{R_1}\frac{\partial^2 C_{R_1}(x,t)}{\partial x^2} + k_1 C_{I_1'}(x,t) \tag{S6.1}$$

$$\frac{\partial C_{I_1}(x,t)}{\partial t} = D_{I_1}\frac{\partial^2 C_{I_1}(x,t)}{\partial x^2} - k_1 C_{I_1}(x,t) \tag{S6.2}$$

$$\frac{\partial C_{I_2}(x,t)}{\partial t} = D_{I_2}\frac{\partial^2 C_{I_2}(x,t)}{\partial x^2} + k_1 C_{I_1}(x,t) - k_2 C_{I_2}(x,t) + k_1 C_{I_1'}(x,t) \tag{S6.3}$$

$$\frac{\partial C_{I_3}(x,t)}{\partial t} = D_{I_3}\frac{\partial^2 C_{I_3}(x,t)}{\partial x^2} + k_2 C_{I_2}(x,t) - 2k_{on} C_{I_3}^2(x,t) \tag{S6.4}$$

$$\frac{\partial C_P(x,t)}{\partial t} = D_P\frac{\partial^2 C_P(x,t)}{\partial x^2} + k_{on} C_{I_3}^2(x,t) \tag{S6.5}$$

$$\frac{\partial C_{R_2}(x,t)}{\partial t} = D_{R_2}\frac{\partial^2 C_{R_2}(x,t)}{\partial x^2} \tag{S6.6}$$

$$\frac{\partial C_{I_1'}(x,t)}{\partial t} = D_{I_1'}\frac{\partial^2 C_{I_1'}(x,t)}{\partial x^2} - k_1 C_{I_1'}(x,t) \tag{S6.7 }$$

The initial concentrations of each species can be expressed as Eq. (S6.8) – (S6.14), where *a* represents the fraction of $OCP^O$ dimer to the total concentration of the reactants.

$$\Delta C_{R_1}(x,0) = -(1-a) \times C_0(1-\cos(qx)) \tag{S6.8}$$

$$\Delta C_{I_1}(x,0) = (1-a) \times C_0(1-\cos(qx)) \tag{S6.9}$$

$$\Delta C_{I_2}(x,0) = 0 \tag{S6.10}$$

$$\Delta C_{I_3}(x,0) = 0 \tag{S6.11}$$

$$\Delta C_P(x,0) = 0 \tag{S6.12}$$

$$\Delta C_{R_2}(x,0) = -a \times C_0(1-\cos(qx)) \tag{S6.13}$$

$$\Delta C_{I_1}(x,0) = a \times C_0(1-\cos(qx)) \tag{S6.14}$$

The calculation procedures are the same as that for OCP(R27L), and the diffusion coefficients and refractive indices for $R_1$, $I_1$, $I_2$, $I_3$ and P was fixed to the values obtained in the calculation for OCP(R27L). As a result of calculation, the signals were well-reproduced with the parameters in Table 2.

**S7. SEC and SAXS measurements of apo-OCP**

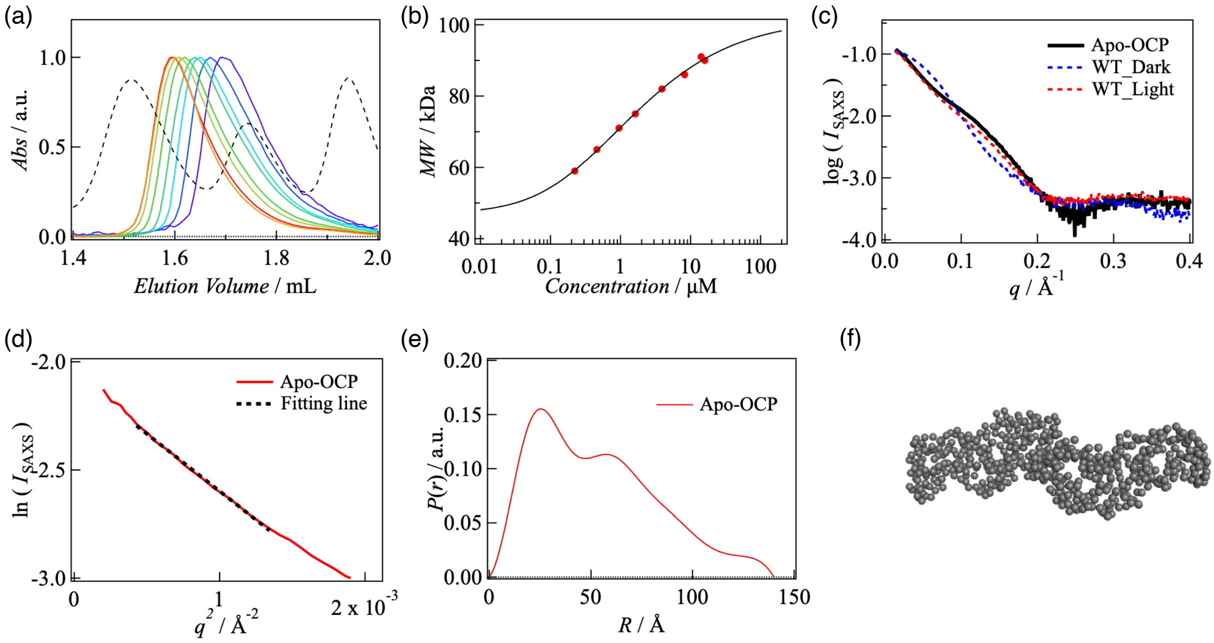


Figure S7. (a) Concentration dependence of the SEC profile for apo-OCP, monitored at 280 nm, with loaded sample concentrations ranging from 300 μM (red) to 2 μM (purple). (b) Plots of molecular mass, determined from the SEC profile, against the concentration at the elution peak. (c) Comparison of SAXS profiles between apo-OCP and holo-OCP. Black solid line: apo-OCP; blue dashed line: holo-OCP (dark state); red dashed line: holo-OCP (light state). (d) Guinier analysis of the SAXS profile for apo-OCP. (e) Radial distribution function obtained from the SAXS profile of apo-OCP. (f) Predicted solution structure of the apo-OCP dimer.
